# New functional vessels form after spinal cord injury in zebrafish

**DOI:** 10.1101/2022.06.09.495446

**Authors:** Ana Ribeiro, Mariana Rebocho da Costa, Carmen de Sena-Tomás, Elsa Charas Rodrigues, Raquel Quitéria, Tiago Maçarico, Susana Constantino Rosa Santos, Leonor Saúde

## Abstract

The vascular system is inefficiently repaired after spinal cord injury in mammals, resulting in secondary tissue damage and immune deregulation that contribute to the limited functional recovery. Unlike mammals, zebrafish can repair the spinal cord and restore motility, but the vascular response to injury has not been investigated. Here we describe the zebrafish spinal cord vasculature, from the body size-dependent vessel ingression during development to the stereotypic vessel organization and barrier specialisation in adulthood. After injury, vessels rapidly regrow into the lesion, preceding the glial bridge and regenerating axons. The initial vascularisation of the injured tissue is done by dysmorphic and leaky vessels. Dysfunctional vessels are later removed, as pericytes are recruited and the blood-spinal cord barrier is re-established. Vascular repair involves an early burst of angiogenesis, likely in response to pro-angiogenic factors detected in the injured spinal cord, including the Vegf pathway. However, the inhibition of the Vegfr2 using genetic and pharmacological methods was not able to efficiently block the formation of new blood vessels, suggesting that other signalling pathways are also involved in this process. This study demonstrates that zebrafish can successfully re-vascularise the spinal tissue, reinforcing the value of this organism as a regenerative model for spinal cord injury.

## Introduction

Spinal cord injury (SCI) causes neuronal and glial loss and severs axonal tracts and blood vessels, leading to severe and permanent consequences for motor and sensory function in mammals (Ahuja et al., 2017). The impact of SCI on the vascular system has both immediate and secondary consequences that worsen the injury outcome. The initial disruption of the vasculature leads to haemorrhage that causes further cell death and allows the extravasation of immune cells (Oudega, 2012; Losey et al., 2014). The Blood-Spinal Cord Barrier (BSCB) is disrupted within minutes of the injury (Whetstone et al., 2003) and can remain compromised for up to 10 weeks (Matsushita et al., 2015). After SCI the vascular system is not correctly repaired, resulting in further leakage and ischemia. Although some blood vessels grow into the lesioned tissue, these vessels have a compromised BSCB due to the disruption of tight junctions (Benton et al., 2008) and incomplete coverage by astrocytes (Casella et al., 2002; Imperato-Kalmar et al., 1997). As new blood vessels enter the lesion site, pericytes also detach from endothelial cells and form the scar tissue (Göritz et al., 2011). The immature vessels continue to leak toxic molecules and inflammatory cells into the tissue, thus contributing to the secondary damage (Hsu et al., 1985; Benton et al., 2008; Takigawa et al., 2010). The impaired vascularisation in the damaged tissue has also been associated to cavitation (Surey et al., 2014).

The improvement of vascular repair could help limit the inflammatory response and the expansion of cystic cavities, as well as support the survival of new and spared cells. Therapies that promote spinal cord re-vascularisation have been explored, but with mixed results (Oudega, 2012; Yao et al., 2021). The therapeutic induction of angiogenesis without additional stabilisation leads to the formation of dysfunctional vessels and worsens the injury outcome (Benton et al., 2003). Consistent with this hypothesis, the combination of VEGF and bFGF (angiogenesis) with Ang-1 (vessel stabilisation) has shown more promising results (Yu et al., 2016). Likewise, therapeutic approaches promoting the recovery of the BSCB also led to some improvement of spinal cord function (Kumar et al., 2017). Biomaterial application and cell transplantation interventions have also been used to promote angiogenesis and BSCB recovery after SCI (Rocha et al., 2018; Yao et al., 2021). However, further work is needed to find more efficient strategies to promote vascular repair while also allowing functional recovery.

The study of vertebrate models that can regenerate the spinal cord, such as zebrafish, could help design re-vascularisation strategies to apply in the mammalian spinal cord. Unlike mammals, adult zebrafish are able to fully restore their motor function in 4 to 6 weeks after SCI (Becker et al., 2004; Hui et al., 2010). The mechanisms underlying the successful regenerative program in zebrafish include the quick resolution of inflammation and absence of a glial scar (Hui et al., 2010); formation of new neurons that replace lost cells (Reimer et al., 2008); and axonal regeneration across the injury site and reconnection to targets (Becker et al., 1997).

However, little is known about the spinal cord vasculature in adult zebrafish and how blood vessels respond to SCI. Brief descriptions of vessel distribution in the adult zebrafish spinal cord in previous studies show blood vessels in the proximity of the central canal (Fang et al., 2014; Liu et al., 2016; Matsuoka et al., 2017) with an enrichement in the grey matter (Mollmert et al., 2020), and increased vessel density after SCI (Liu et al., 2016).

Here, we carried out a comprehensive analysis of vascular distribution and morphology during zebrafish spinal cord development, homeostasis and regeneration. The stereotypic organisation and permeability properties of the spinal vascular network were disrupted by SCI. But unlike mammals, the zebrafish spinal cord was able to re-establish a stable and functional vascular network, driven by endothelial proliferation, pericyte recruitment and BSCB recovery. Re-vascularisation was only partially dependent on Vegfr2 signalling, suggesting other pro-angiogenic pathways are active.

## Results

### Spinal cord vascularisation in developing zebrafish

The intraspinal tissue is vascularised only late in development, as vessels grow from the perineural vascular plexus (PNVP) that surrounds the spinal cord and enter dorso-laterally into the neural tissue (Matsuoka et al., 2017). This process was described to occur between 12-14 (Wild et al., 2017) or 17-18 days post fertilisation (dpf) (Matsuoka et al., 2017), suggesting that the timing of the entry cannot be predicted by age alone. Our detailed characterisation of spinal vessel ingression also revealed that it could occur in a range of ages and varied with the body region analysed. Moreover, we observed that body length at 6 weeks post fertilisation (wpf) could range from 4.7 mm to 10 mm and were undistinguishable from fish one week younger (5 wpf) (Fig. 1A,B). This observation led us to investigate which parameters, other than age, were better correlated with the entry of vessels into the spinal cord. For this we obtained sections of the trunk region of fish with 5 and 6 wpf and quantified the number of vessels (labelled with the endothelial reporter *Tg(kdrl:ras-mCherry)*^*s896*^) per section at different positions along the trunk region (Fig. 1A,C,D). In addition, we measured the area, left-right (LR) length and dorsal-ventral (DV) length of the spinal cord in each section (Fig. 1E). We performed a correlation matrix analysis between all the parameters quantified (Fig. 1F). As expected, the parameters related with size (spinal cord area, DV length, body length and LR length) showed a high positive correlation between each other, a low correlation with age and a negative correlation with rostral-caudal (RC) position, since the spinal cord size decreases along the RC level. The number of vessels also showed a high positive correlation with size-related parameters and low correlation with age, suggesting that the size of the fish and, in association, the size of the spinal cord, have the strongest influence in the timing of intraspinal vascularisation. Likewise, the number of vessels was negatively correlated with RC position, consistent with a rostral to caudal pattern of vessel ingression. The parameter best correlated with the number of vessels was spinal cord area. Blood vessels started entering the spinal cord trunk region in 6 mm fish, when the spinal cord area reached an average of 8378 ± 2106 µm^2^ (Fig. 1G). This area corresponded to an approximate radius of 50 µm, suggesting that below this radius the PNVP is sufficient to perfuse the intraspinal tissue and above this threshold vessels start invading the spinal cord. At this stage most cells in the spinal cord are differentiated neurons (HuC/D^+^) and only a few progenitor cells remain around the central canal (Fig. 1C,D), unlike the more immature cell composition observed in mouse and chicken at the stage of vessel entry (Himmels et al., 2017). These data describe the size-dependent vascularisation of the zebrafish spinal cord at a later developmental stage than in mammals, possibly coordinated by different cell types.

**Figure 1.**
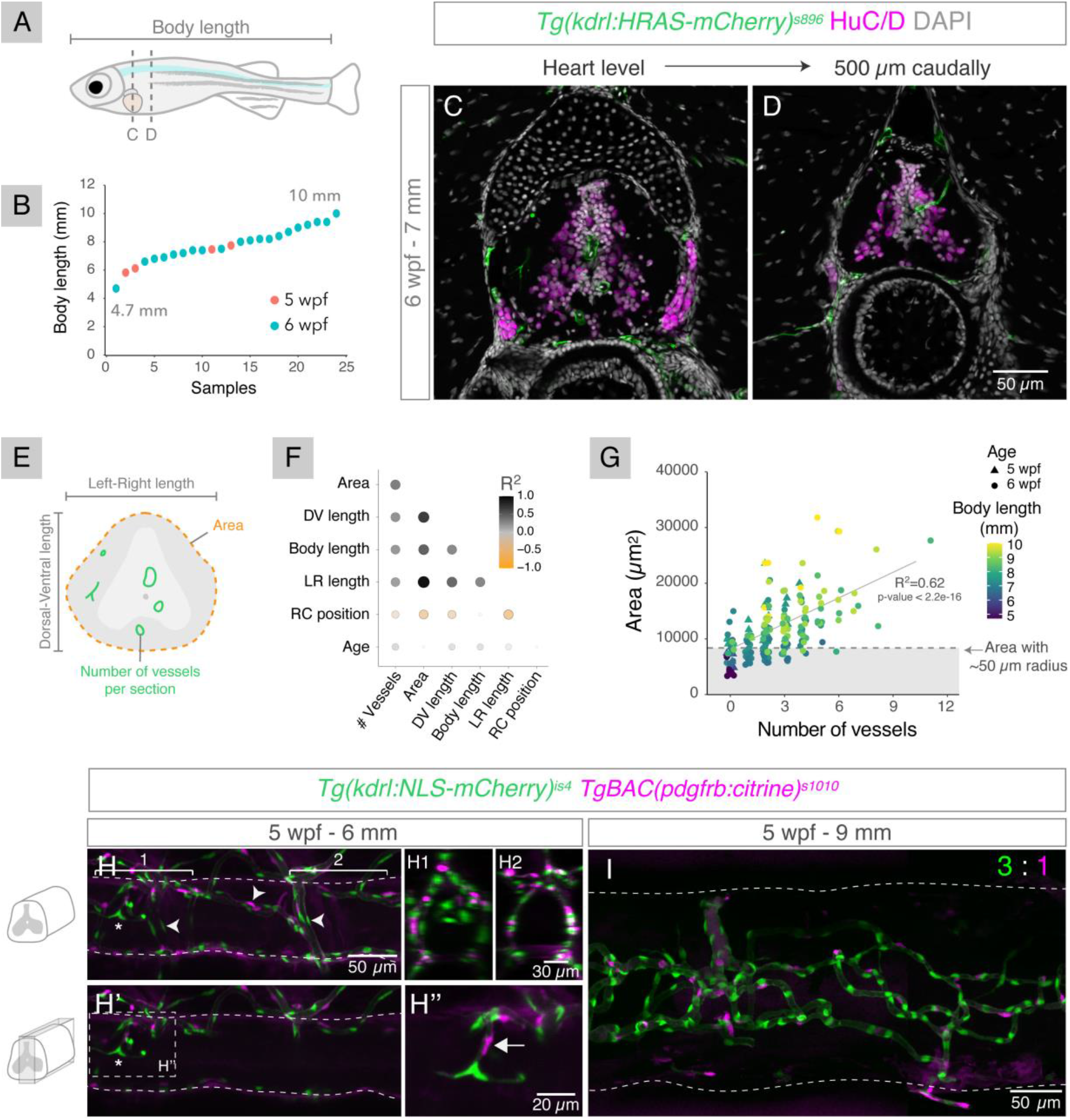
Spinal cord vascularisation in juvenile zebrafish. A. Definition of body length and rostral-caudal region analysed in juvenile zebrafish. B. Body length in individual samples with 5 or 6 weeks post-fertilisation (wpf). C,D. Confocal projections of sections of 6 wpf fish at the heart level (C) and 500 µm caudally (D) with labelled endothelial cells (*Tg(kdrl:ras-mCherry)*^*s896*^) and co-labelled for HuC/D^+^ neurons and nuclear DAPI staining. The rostral region shows more vessels than the caudal region. E. Schematic showing the parameters quantified in the imaged sections. F. Correlation matrix dot plot of the quantified parameters: area; DV (dorsal-ventral) length; body length; LR (left-right) length; RC (rostral-caudal) position (starting at the heart level); age and number of vessels per section. Dot size and colour represent the Spearman correlation coefficient. G. Spinal cord area relative to number of vessels per section in all sections analysed. Symbol shape indicates age and symbol colour indicates body length. Dashed line corresponds to average spinal cord area in sections with 1 vessel inside. H-I. Confocal images of 5 wpf juveniles with different sizes with endothelial nuclei in green (*Tg(kdrl:NLS-mCherry)*^*is4*^) and pericytes in magenta (*TgBAC(pdgfrb:citrine)*^*s1010*^), in a projection of the whole spinal cord region (H,I) or just the central region (H’). In H-H’, Vessels around the spinal cord (perineural vascular plexus) are highlighted with arrowheads and a vessel inside the spinal cord is identified with an asterisk. Transversal views of the image H show a region with vessels inside (H1) and a region empty of vessels (H2). A magnified view of the image H’ shows a pericyte (arrow) co-recruited with an ingressing blood vessel (H’’). In larger fish (I) the spinal cord vasculature is more complex and shows a higher coverage by pericytes (3 endothelial cells per pericyte, n=3).

The mature spinal cord microvasculature is coated by perivascular cells - pericytes - that contribute to the maintenance of the BSCB (Winkler et al., 2012), including in zebrafish (Tsata et al., 2021). To determine if pericytes also accompany endothelial cells (ECs) as they enter the spinal cord, we analysed 5 wpf fish with labelled pericytes (*TgBAC(pdgfrb:citrine)*^*s1010*^) and ECs (*Tg(kdrl:NLS-mCherry)*^*is4*^), using confocal microscopy to image cleared wholemount fish (in smaller samples) or dissected spinal cords. We observed pericytes being co-recruited with the angiogenic sprouts (Fig. 1H-H’’) as ECs invade the nervous tissue (in 6 mm fish), and a high density of pericyte coverage (1 pericyte to 3 ECs) (Fig. 1I) when a more developed vascular network was present (in 9 mm fish). The vessels around the spinal cord (PNVP) are also surrounded by pericytes (Fig. 1H; Tsata et al., 2021), raising the possibility that the PNVP-associated pericytes are the source of intraspinal pericytes. These results show the coordinated entry and dispersal of ECs and pericytes during the formation of the spinal cord vasculature.

### Spinal cord vasculature in adult zebrafish

We next characterised the spinal cord vasculature in the adult zebrafish. We used light sheet microscopy to acquire wholemount images of cleared spinal cords of transgenic fish with fluorescently labelled ECs (Fig. 2A,B). This analysis revealed a stereotypic organization of blood vessels, with preferential distribution in the grey matter (Fig. 2B’). Ingression points at dorsal and ventral positions connected the external plexus to the internal vasculature. The ventral vessels entered towards the centre of the tissue and ran along the central canal with some branching (Fig. 2B1,B2). The dorsal vessels ramified into a more complex capillary network that spread into the central and lateral tissue (Fig. 2B2,B3), mostly in close proximity to neurons (Fig. 2C). This distribution suggested that the zebrafish spinal cord was only supplied by a central system and lacked the peripheral system (vasocorona) present in humans (Bosmia et al., 2015). To determine if these peripheral vessels were being missed due to the endothelial reporter used (*kdrl/vegfr2*), we analysed the distribution of a broader endothelial marker, *fli1* (Küchler et al., 2006, Yaniv et al., 2006). The combination of a *fli1* reporter (*Tg(fli1:EGFP)*^*y1*^) with a *kdrl* reporter indeed revealed single positive *fli1* vessels, but these were restricted to the exterior of the spinal cord (Fig. 2D,D’) and were possibly lymphatic vessels (Chen et al., 2019). These findings support the absence of a vasocorona in the zebrafish spinal cord.

**Figure 2.**
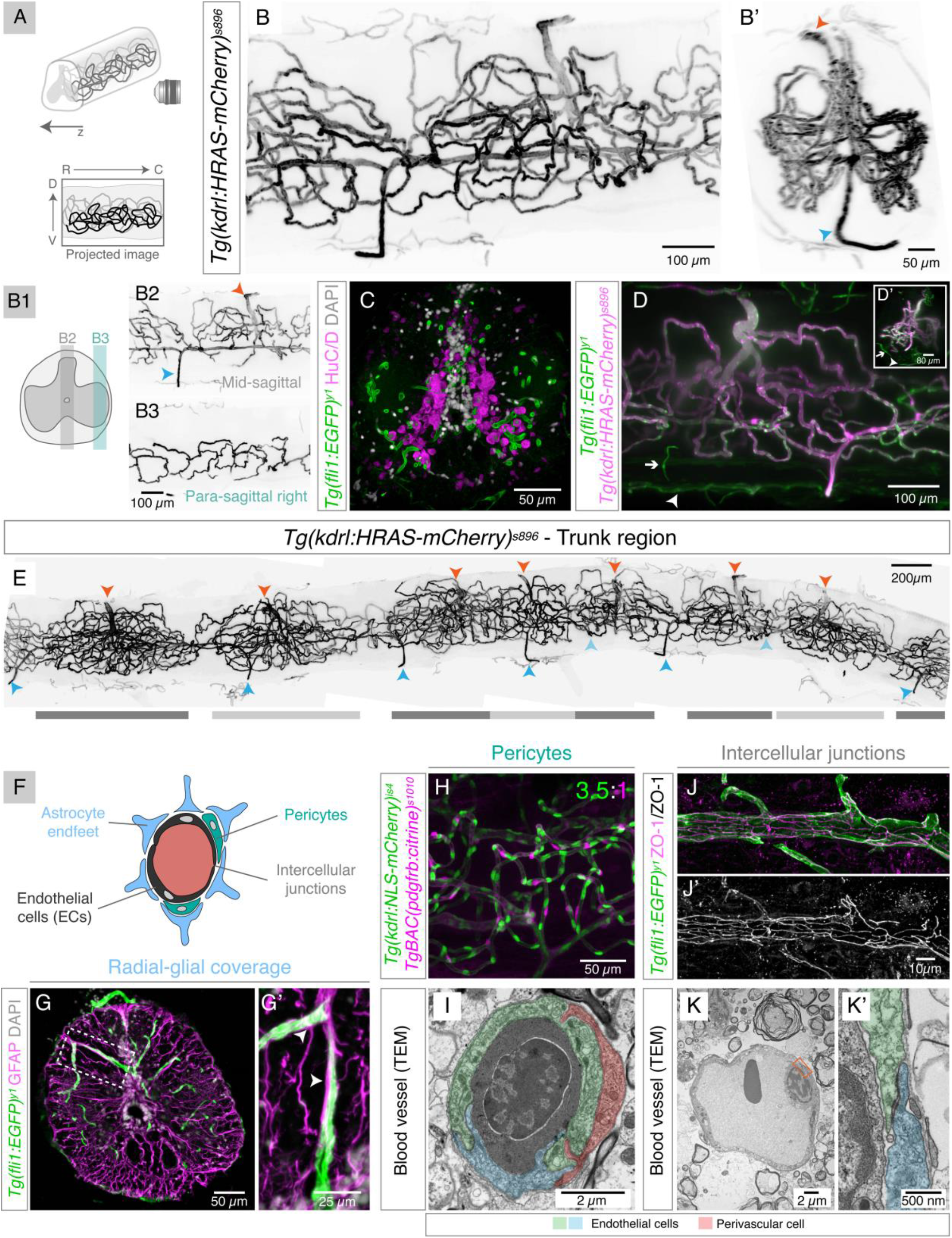
Organization of the spinal cord vasculature in adult zebrafish. A. Schematic of light sheet image acquisition of wholemount cleared spinal cords, with the z axis from the right to left. Below, a schematic of the projected light sheet image shows the rostral (R) to caudal (C) and ventral (V) to dorsal (D) orientation. B. Projection of the whole spinal cord with labelled endothelial cells (ECs) (*Tg(kdrl:ras-mCherry)*^*s896*^). A transversal view of the image (B’) shows vessels ingressing in the dorsal (orange arrowhead) and ventral (blue arrowhead) regions and dispersing mostly into the grey matter. B1. Schematic of a transverse spinal cord depicting the 100 µm regions projected in B2,B3. Dorsal and ventral ingressing vessels (arrowheads) branch into the central (B2) and lateral (B3) regions. C. Image of transverse section of *Tg(fli1:EGFP)*^*y1*^ spinal cords co-labelled for HuC/D^+^ neurons and nuclear DAPI staining, showing blood vessels in close proximity to neurons. D. Projection of the whole spinal cord with *fli1*^*+*^*/kdrl*^*+*^ intrinsic vessels and *fli1*^*+*^*/kdrl*^*-*^ extrinsic vessels surrounding the spinal cord (arrows). Inset with transversal view of the image (D’) E. Composite of the trunk spinal cord vasculature (*Tg(kdrl:ras-mCherry)*^*s896*^) in a 4.6 mm region. Arrowheads label intraspinal vessels connecting to the outside vasculature in ventral (blue) and dorsal (orange) regions. Predicted positions of some of the ventral vessels are shown in light blue. Dark and light grey bars below image identify putative segments of the vasculature. F. Schematic of the blood-spinal cord barrier components. G. Confocal projection of a transverse section of a *Tg(fli1:EGFP)*^*y1*^ spinal cord with ECs in green and co-labelled with an anti-GFAP antibody in magenta. G’. Magnification of box in (G). GFAP labels a population of radial glial cells that are in contact with ECs (arrowheads). H. Confocal projection of wholemount spinal cord with endothelial nuclei in green (*Tg(kdrl:NLS-mCherry)*^*is4*^) and pericytes in magenta (*TgBAC(pdgfrb:citrine)*^*s1010*^). The adult zebrafish spinal cord has an average of 3.5 ECs per pericyte. I. TEM image of a cross-section of a blood vessel, showing two contacting ECs (in blue and green) and an associated pericyte (in red). J,J’. Confocal image of a *Tg(fli1:EGFP)*^*y1*^ spinal cord with ECs in green, co-stained with the tight junction marker ZO-1 in magenta. Tight junctions are detected in blood vessels. K. Transmission Electron Microscopy (TEM) image of a blood vessel, with a magnification of the orange box in (K’) showing intercellular junctions between ECs.

A wider perspective of the spinal cord revealed a segmented vasculature, with clusters of vessels separated by less vascularised regions (Fig. 2E). The complexity and distributions of the segments was not homogenous along the spinal cord and may reflect different metabolic requirements of the spinal tissue along the rostral-caudal axis.

The central nervous system microvasculature displays specialised modifications that control the transport of compounds and cells into the tissue: ECS are sealed by tight junctions and have a selective transport system; the endothelial layer is covered by pericytes, enveloped in a basal lamina and wrapped by astrocyte endfeet (Profaci et al., 2020) (Fig. 2F). We examined the presence of BSCB components using immunohistochemistry to detect GFAP^+^ glial cells and *Tg(pdgfrß:citrine)*^*s1010*^ to label pericytes. GFAP+ projections were detected adjacent to ECs, indicating that glial cells encircle blood vessels (Fig. 2G,G’)). Consistent with a previous study (Tsata et al. 2021), pericytes were detected in close association to ECs (Fig. 2H). The quantification of the number of pericytes and ECs showed a high coverage by pericytes, with a ratio of 1 pericyte per 3.5±0.6 ECs. This ratio is close to that observed in the CNS of amniotes (around 1:1) (Shepro & Morel, 1993). Perivascular cells were also visualised in Transmission Electron Microscopy (TEM) images, which showed peg-socket junctions between pericytes and ECs (Fig. 2I). Blood vessels displayed tight junctions (identified using tight junctions marker ZO-1) (Fig. 2J,J’) and also adherent junctions, desmosomes and gap junctions (detected by TEM) (Fig. 2K,K’). This characterisation reveals that, as in mammals, the adult zebrafish spinal cord displays blood vessels with specialised modifications associated with a mature BSCB.

### Blood vessels rapidly re-vascularise the injured spinal cord

To examine how blood vessels respond to a SCI, we performed a contusion injury model in adult zebrafish (Hui et al., 2010) and imaged the blood vessels in cleared wholemount samples (*Tg(kdrl:ras-mCherry)*^*s896*^) at different days post injury (dpi). To quantify vessel length and morphology we used a 3D image analysis workflow developed by Kugler et al. (Kugler et al., 2022), which measures vessel branch length and position, thickness and tortuosity (Supp. Fig. 1). At 3 dpi the vasculature was still severed at the injury site, with an overall reduction in vessel length when compared to uninjured spinal cords (Fig. 3A-B’, F). New vessels rapidly re-vascularised the lesioned region and formed agglomerates by 7 dpi, resulting in an increase in vessel length (Fig. 3C,C’, F). In addition, the vessels from the rostral and caudal sides appeared to have anastomosed at the injury centre, pointing to a continuity in blood flow between the two sides of the injury. The vessels clusters were still visible at 14 dpi, but less so at 30 dpi (Fig. 3D-E’). While total vessel length remained constant from 7 dpi (Fig. 3F), vessel density was not homogeneous in all domains. Along the rostral-caudal axis, the injury region failed to recover to pre-injury levels and the rostral side showed a better recovery than the caudal side (Fig. 3G, top panel). Along the dorsal-ventral axis, the recovery was lowest in the middle region, where vessel density had been highest pre-injury, and the ventral region improved more than the dorsal region (Fig. 3G, middle panel). Along the left-right axis (z axis) the uninjured spinal cord showed a higher accumulation of vessels in the central region, in proximity of the central canal, than in the lateral regions (data not shown). The central region showed a better recovery after an injury than the lateral region, indicating that new vessels grow preferentially in the central canal area (Fig. 3G, bottom panel).

**Figure 3.**
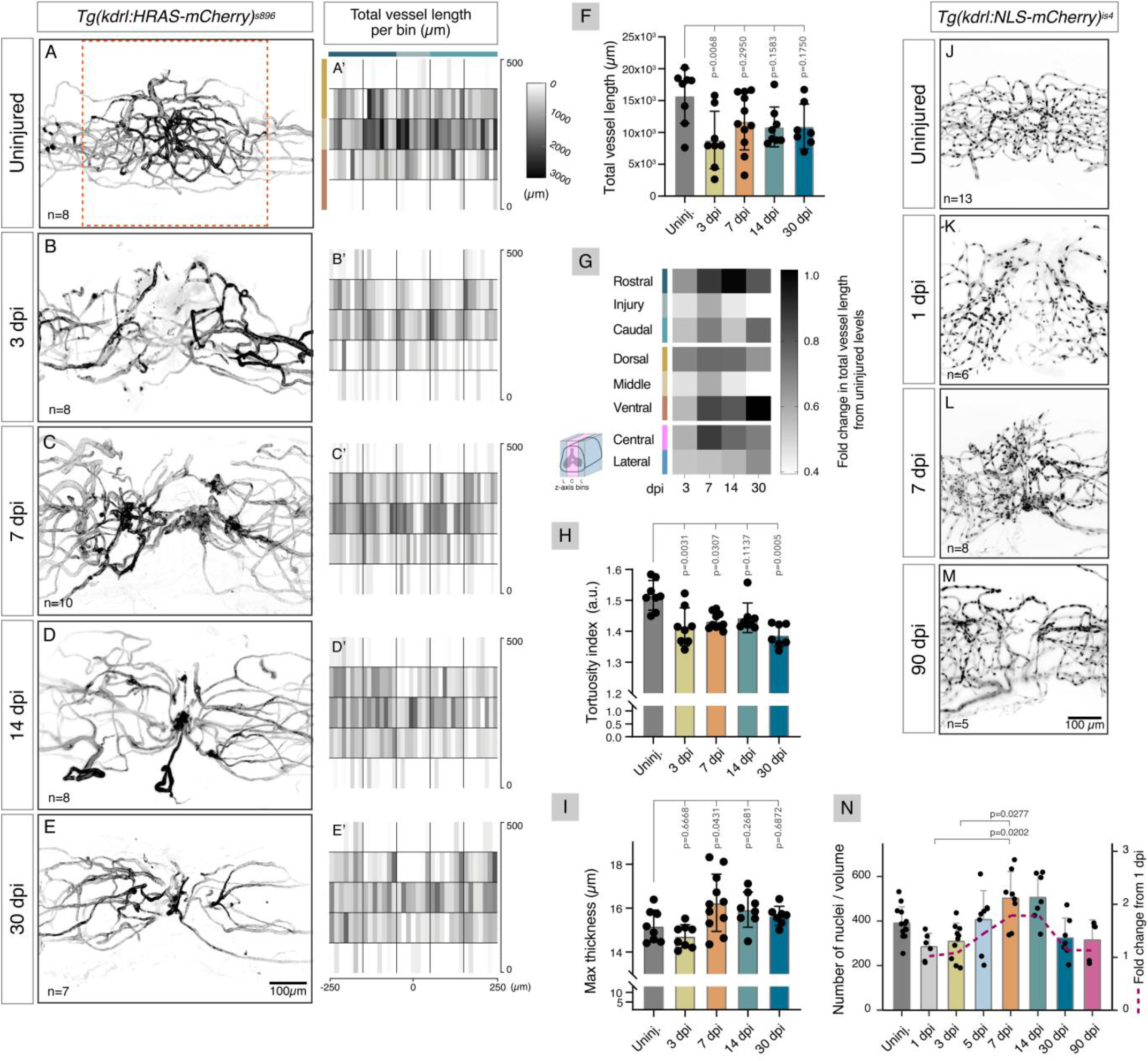
Changes in vascular distribution and morphology in response to spinal cord injury. A-E. Projections of light sheet microscopy images of whole spinal cords with labelled endothelial cells (*Tg(kdrl:ras-mCherry)*^*s896*^) in uninjured and 3, 7, 14 and 30 days post-injury (dpi) with contusion. The vasculature is disrupted at 3 dpi, but new vessels rapidly appear at 7 dpi and some are removed/remodelled by 30 dpi. A’-E’. Heatmap of the total vessel length per 100 µm bins along the rostro-caudal and dorso-ventral axis in a 500×500 µm region (orange square in A). Each rectangle inside a bin corresponds to an individual sample (n=7-10). F. Average vessel length over time post-injury. G. Fold change in total vessel length relative to uninjured levels over time post-injury in different regions along the rostral-caudal axis, the dorsal-ventral axis and the left-right axis. H. Average vessel tortuosity index (vessel length / euclidean distance) over time post-injury shows more linear vessels are formed after injury. I. Maximum vessel thickness over time post-injury shows vessels with larger caliber are present in the repaired vasculature. J-M. Projections of light sheet images of whole spinal cords with labelled endothelial nuclei (*Tg(kdrl:NLS-mCherry)*^*is4*^), without injury and 1, 7 and 90 days post-injury (dpi) with contusion. N. Number of endothelial nuclei over time post-injury in a 500×500 µm region of the projected images. The blue dashed line indicates the fold change in the number of nuclei relative to 1 dpi (right side axis). Data represent means ± SD. Statistical tests: Kruskal-Wallis test followed by Dunn’s multiple comparisons post-hoc test relative to uninjured control (F,H,N); One-way ANOVA followed by Dunnett’s multiple comparisons test (I) (p-values > 0.05 in (N) are not shown).

We also examined the morphology of the vessels, namely tortuosity and vessel calibre. Blood vessels in the injured spinal cord showed a lower tortuosity index than vessels in the uninjured spinal cord (Fig. 3H), indicating that vessels formed after an injury are overall more linear than the normal vasculature. However, some sites of vascular agglomeration found at 7 and 14 dpi displayed very tortuous vessels (Fig. 3C, D). In addition, vessels with a larger calibre than normal were detected at 7 and 14 dpi (Fig. 3I).

We also investigated if the type of injury influenced the vascular response. We compared contusion and transection (Becker et al., 1997) injury models and observed similar vascular dynamics in both models (Supp. Fig. 2A-F). Using the transection model, in which all axons and glia are eliminated in the injury site and new axons and glial projections are more easily identified (Supp. Fig. 2G,H), we examined how re-vascularisation is coordinated with glial bridge formation (*Tg(gfap:GFP)*^*mi2001*^) and axonal regrowth (acetylated alpha-tubulin) (Goldshmit et al., 2012; Mokalled et al.. 2016). We observed that at 3 dpi, when new axons and glial cells were still absent, vessels were already detected close at the injury site (Supp. Fig. 3A), revealing that blood vessels are among the first structures to enter the lesioned tissue. In addition, at 5 dpi, when axons were observed growing into the injured tissue, some axons were detected near blood vessels, with or without accompanying glial projections (Supp. Fig. 3C). This observation points to a possible early association between vascular and axonal growth.

To further investigate how the vasculature was repaired we quantified the number of ECs in the injured area from 1 to 90 dpi using a nuclear endothelial label (*Tg(kdrl:NLS-mCherry)*^*is4*^). The quantification of endothelial nuclei showed similar dynamics to the quantification of vascular length: the number of ECs decreased after the injury, was followed by a recovery at 7 dpi with the formation of vessel agglomerates and decreased again from 30 dpi onwards (Fig. 3J-M,N, Supp. Fig. 4). The reduction in vascular complexity and cell number observed in later time-points (60 and 90 dpi) suggests that during the rapid repair process vessels are formed in excess and some are dysfunctional and later are removed or remodelled.

In summary, these findings reveal an initial phase of rapid endothelial growth with formation of large and tortuous vessels that agglomerate in the central region, where the presence of progenitor cells and/or surrounding neurons possibly influences the vascular response. The repaired vasculature appears to be optimised later, resulting in a simplified version of the vascular network, with fewer and more linear vessels than in control spinal cords. Critically, many of the newly formed vessels persist at 90 dpi, suggesting that these vessels are functional and provide long-term support to the damaged tissue, unlike what has been described in the mammalian spinal cord (Casella et al., 2002).

### Reestablishment of barrier properties in the repaired vasculature

To determine if the new vessels were functional, i.e., if vessels were perfused and developed a BSCB, we injected the 10KDa dextran Alexa Fluor 647 (A647) into circulation (Fig. 4A). This tracer dye is size-restricted from the CNS (Jeong et al. 2008). To assess perfusion of new vessels we allowed the dye to circulate during 1 minute before collecting the spinal cord. The distribution of dextran-A647 in cleared wholemount spinal cords was analysed using light sheet imaging (Fig. 4B-D’’). We confirmed that the injection was successful by acquiring a region caudal to the injury (Fig. 4B’’-D’’). In this region the dextran-A647 remained inside blood vessels. At 1 dpi the tracer was absent from most vessels in the injured region (Fig. 4B,B’), suggesting that vessels were not yet perfused or the dye was not retained inside the vessels. At 5 dpi, increased dye levels were observed inside vessels, particularly in more lateral regions (Fig. 4C,C’). By 7 dpi most vessels at the injury centre were filled with the tracer (Fig. 4D,D’), indicating that the growing vessels had successfully fused and blood was flowing across the injury site.

**Figure 4.**
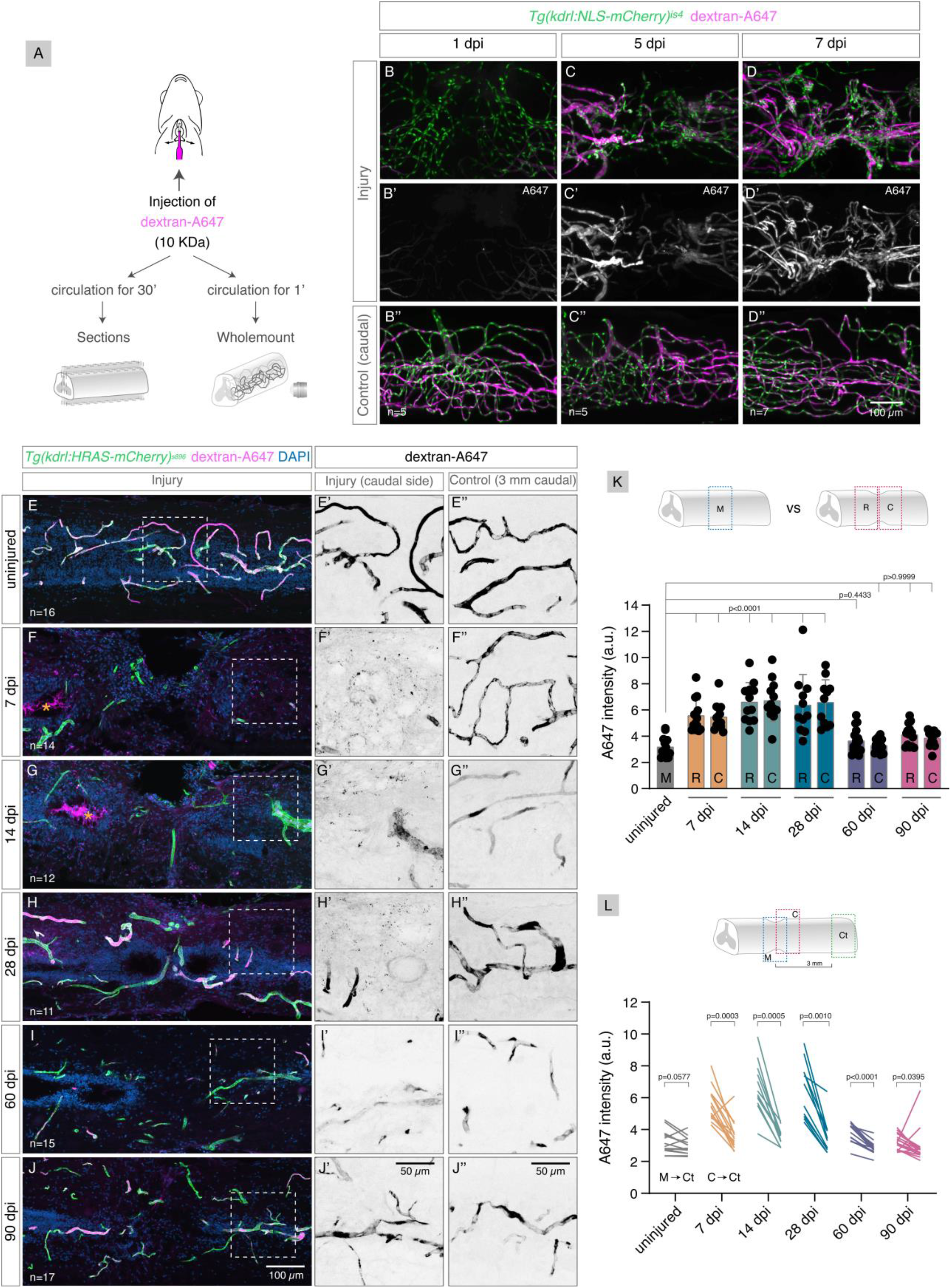
Reestablishment of the blood-spinal cord barrier (BSCB) after injury. A. Schematic of the vessel perfusion/permeability assay, in which a fluorescent dye that is size-restricted by the BSCB (10KDa dextran-Alexa647 (A647)) was injected into the heart and allowed to perfuse for either 1 minute (to measure vessel perfusion) or 30 minutes (to measure vessel leakiness). The spinal cord was then processed for wholemount light sheet imaging (1’) or confocal imaging in sections (30’). B-D’’. Projections of light sheet images of whole spinal cords with labelled endothelial nuclei (*Tg(kdrl:NLS-mCherry)*^*is4*^), perfused with dextran-A647 for 1 minute and analysed at 1, 5 and 7 days post-injury (dpi) with contusion. Vessels in the injury region are perfused around 7 dpi (B’-D’). A region 2 mm caudally to the injury is used as control of the injection (B’’-D’’). E-J. Confocal projections of contused spinal cords longitudinal sections with labelled endothelial cells (*Tg(kdrl:ras-mCherry)*^*s896*^), perfused with dextran-A647 for 30 minutes and with DAPI-stained nuclei. Orange asterisks highlight the accumulation of dextran-A647 in the central canal in F,G. Magnified views of the dextran-A647 signal in the caudal side of the injury are shown in E’-J’. and a control region 3 mm caudal to the injury is shown in E’’-J’’. Speckles of leaked dextran-A647 are detected outside vessels at 7, 14 and 28 dpi (F’,G’,H’). K. Quantification of extravascular intensity of dextran-A647 in the rostral and caudal side of the injury in 7, 14, 28, 60 and 90 dpi spinal cords and compared with the corresponding middle region (M) in uninjured spinal cords. Extravascular dextran-A647 levels only return to uninjured levels at 60 dpi. L. Comparison of dextran-A647 extravascular intensity in the caudal side of the injury (or a corresponding region in uninjured spinal cords) with a control region 3 mm caudal to the injury. Each line shows the change between the 2 regions in individual spinal cords. Extravascular levels in the injured region are significantly different from the control region at 7, 14, 28 and 60 dpi. Data represent means ± SD. Statistical tests: Kruskal-Wallis test followed by Dunn’s multiple comparisons post-hoc test relative to uninjured control (K) and Wilcoxon matched-pairs signed rank test (L).

To determine if the BSCB was re-established in the repaired vasculature we injected the dextran-A647 into circulation and allowed the dye to circulate during 30 minutes before collecting the spinal cord. The dye is expected to remain inside vessels with an intact barrier but will diffuse into the tissue over time if the vessels are leaky (O’Brown et al., 2019). We analysed the distribution of dextran-A647 in sections in uninjured spinal cords and from 7 dpi (when new vessels are perfused) until 90 dpi (Fig. 4E-J). The intensity levels of dextran-A647 were measured in the tissue parenchyma in the rostral and caudal sides in the injury or in control regions (uninjured spinal cord or region distant from the injury) (Supp. Fig. 5A-B’). The image analysis revealed that little dye extravasated in uninjured spinal cords (Fig. 4E,E’) or in control regions 3 mm caudal to the injury (Fig. 4E’’-J’’), indicating that the barrier properties of vessels are intact. By contrast, speckles of dye were detected in the extravascular tissue at 7, 14 and 28 dpi (Fig. F-G,F’-H’), revealing that vessels are still permeable at these time points. This speckled distribution was only present in the injured tissue when dextran-A647 was injected (Supp. Fig. 5C-D’), indicating that the signal is specific for the extravasated dye. The dye was present not only in the parenchyma, but also in the central canal lumen (Fig. 4F,G). The dextran-A647 possibly contaminates the cerebrospinal fluid through the choroid plexus (Daneman et al., 2010). The quantification of A647 intensity in the parenchyma confirmed that the dye levels are significantly increased in the injured region at 7, 14 and 28 dpi when compared to uninjured samples (Fig. 4K) or to a distant region in the same spinal cord (Fig. 4L). Moreover, this quantification revealed that A647 leakage is comparable between the rostral and the caudal sides of the injury. By 60 dpi, the parenchymal A647 levels had returned to uninjured levels and remained so at 90 dpi (Fig. 4I-J’’,K), although A647 levels in the injured tissue were still above caudal levels (Fig. 4L). This indicates that the barrier properties of the repaired vasculature are re-established between 28 and 60 dpi. These results reveal that the BSCB development is a delayed process when compared with new vessel formation, suggesting that the spinal cord tissue remains exposed to circulating compounds and cells for a prolonged period.

### Angiogenesis contributes to new vessel formation

To investigate how the vasculature is repaired, we focused on the phase of new vessel formation. From 1 to 7 dpi, the number of ECs showed an almost 2-fold increase (from 284.5 ± 60.4 to 502.6 ± 120.3) (Fig. 3N). This fold change suggests that endothelial proliferation and/or migration contribute to the regrowth of vessels into the injured tissue. To assess the contribution of endothelial proliferation to the formation of new blood vessels we performed EdU incorporation after spinal cord/sham injury. To evaluate the cumulative proliferation of ECs, we injected EdU in 3 consecutive days (2-4 dpi) and collected the spinal cords at 5 dpi (Fig. 5A-B’’). The quantification of EdU+ ECs showed a significant increase in the number of proliferating ECs in injured spinal cords, when compared to sham injuries (Fig. 5F). In fractional terms, this represented an increase of accumulated proliferative ECs relative to the total number of ECs from an average of 0.2% ± 1.0 in sham animals to 15% ± 7.7 of ECs in injured animals (Fig. 5G). These results indicate that endothelial proliferation contributes to the repair of damaged blood vessels.

**Figure 5.**
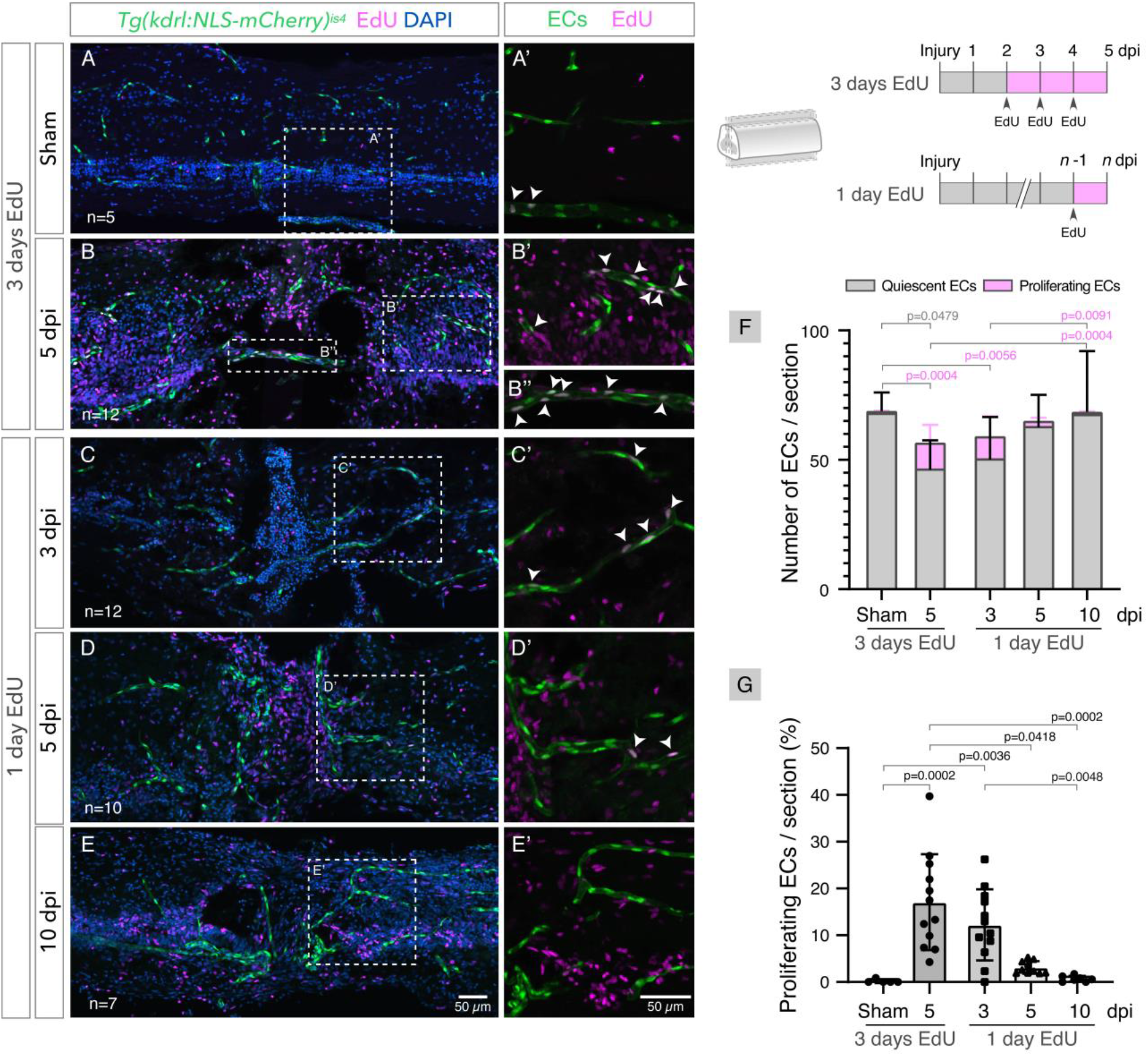
Endothelial proliferation during vascular repair. A-E. Confocal projections of longitudinal sections of contused spinal cords with labelled endothelial cells (*Tg(kdrl:NLS-mCherry)*^*is4*^), EdU staining and nuclear DAPI staining, with magnified views (A’-E’) showing proliferating ECs (arrowheads). Repeated EdU injections were performed 3 consecutive days before spinal cord collection at 5 days after sham/spinal cord injury (A,B). Single EdU injection were administered 1 day before spinal cord collection at 3, 5 and 10 dpi (C,D,E). F. Total number of endothelial cells (ECs) per section, grouped as proliferating (EdU^+^) or quiescent (EdU^-^) cells. G. Fraction of proliferating ECs (EdU^+^ ECs / total ECs) per section. Data represent means ± SD. Statistical tests: Kruskal-Wallis test followed by Dunn’s multiple comparisons post-hoc test (F,G) (p-values > 0.05 are not shown).

To examine the dynamics of EC proliferation in more detail, we performed single-pulse EdU incorporation one day before spinal cord collection at 3, 5 and 10 dpi (Fig. 5C-E’). The quantification of EdU+ ECs showed the highest number of proliferating ECs at 3 dpi and gradually decreased until 10 dpi (Fig. 5F)). In terms of the fraction of ECs that proliferated after an injury, an average of 11.2% ± 11.6 of ECs were proliferating at 3 dpi, while only 3% ± 3.8 and 0.6% ± 1.9 of ECs proliferated at 5 and 10 dpi, respectively (Fig. 5G). These data reveal that an early and transient proliferative response contributes to the repair of the injured vasculature.

### Pericyte recruitment during vessel stabilisation

During the repair of a disrupted vasculature, the formation of new vessels needs to be coordinated with the recruitment of support cells, including pericytes. To determine if and when pericytes were recruited to the vessels, we analysed transgenic zebrafish with labelled pericytes and endothelial nuclei (*Tg(pdgfrß:citrine)*^*s1010*^; *Tg(kdrl:NLS-mCherry)*^*is4*^) up to 90 days after a spinal cord contusion/sham injury. We observed a higher density of pericytes surrounding repairing vessels, with a gradual increase from 3 dpi until 30 dpi, with significantly higher numbers from 7 to 30 dpi when compared to a sham injury (Fig. 6A-H,I). Pericyte proliferation, detected in early time points, contributed to the increase in pericyte number (Supp. Fig. 6). From 30 dpi the number of pericytes decreased until the last time point analysed, 90 dpi, when pericytes reached pre-injury levels. A similar dynamic was observed for the number of ECs (Fig. 6I). To determine if the number of pericytes changed proportionally to the number of ECs, we calculated the ratio of ECs:pericytes, in which a low ratio represents a high coverage of blood vessels by pericytes (Fig. 6J). This analysis showed an increased pericyte coverage in all time-points post-injury when compared to sham injury. The highest pericyte coverage was observed at 30 and 60 dpi, which also corresponds to the period when the BSCB is being re-established. By 90 dpi, when the barrier has recovered, pericyte coverage was returning to homeostatic levels. The correlation between peak pericyte coverage and BSCB repair points to a possible involvement of pericytes in the reestablishment of the barrier.

**Figure 6.**
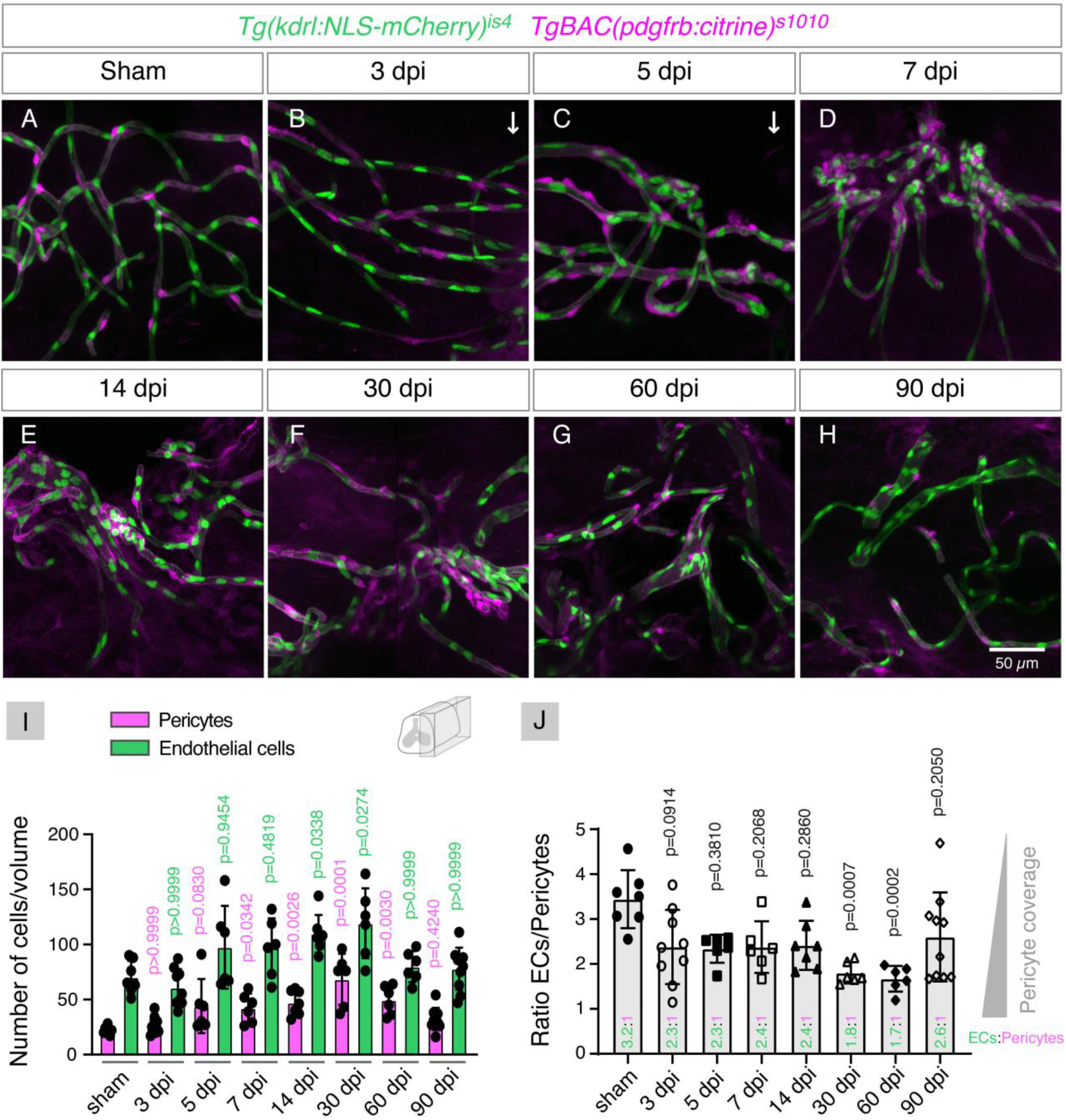
Pericyte recruitment during vascular repair. A-H. Confocal projections of the lateral region (90 µm depth) of wholemount spinal cords with labelled endothelial nuclei (*Tg(kdrl:NLS-mCherry)*^*is4*^) and pericytes (*TgBAC(pdgfrb:citrine)*^*s1010*^) after sham injury and between 3 and 90 dpi. The images show the centre of the injury, except in B,C, where an arrow identifies the position of the injury. I. Total number of pericytes and ECs over time post-injury. J. Ratio of ECs per pericyte over time post-injury. The value of the average ratio for each time-point is shown at the base of the bar. Lower ratios correspond to higher coverage of vessels by pericytes. Data represent means ± SD. Statistical tests: Kruskal-Wallis test followed by Dunn’s multiple comparisons post-hoc test relative to uninjured control (I,J).

We also observed that the majority of cells labelled by the *pdgfrb:citrine* transgene were associated with vessels and we were unable to identify a separate population of *pdgfrb*^+^ cells detached from vessels present in the injury core, as previously observed in the mammalian spinal cord (Göritz et al., 2011) and in the larval zebrafish spinal cord (Tsata et al., 2021). However, it is possible that this population is present in the adult zebrafish, but we are unable to detect it due to the high background of the reporter line used (Supp. Fig. 7).

These data show that new vessels recruit an excess of pericytes over a long period. After the remodelling and maturation of the new vasculature, the coating of vessels by pericytes returns to the homeostatic density.

### Formation of new blood vessels depends partially on the Vegfr2/Vegf pathway

To help identify the signalling pathways involved in the formation of new vessels we used qPCR to analyse the expression of genes involved in endothelial regulation and pericyte recruitment in contused spinal cords (Fig. 7A). At 3 dpi, the peak time of endothelial proliferation, the only pro-angiogenic gene showing a small increase was *vegfaa*. At 7 dpi, when blood vessels reach a peak in length and EC number, more genes associated with blood vessel formation were up-regulated (*vegfaa, apln, shha, tie2/angiopoietins angpt1* and *angpt2b*). In addition, genes involved in pericyte recruitment were also up-regulated at 7 dpi, particularly *pdgfbb* and *pdgfrb*. The observed changes in expression point to possible pathways involved in vascular re-growth, while the activation of the Pdgf pathway is consistent with the significant increase in pericyte number in the repaired vasculature.

**Figure 7.**
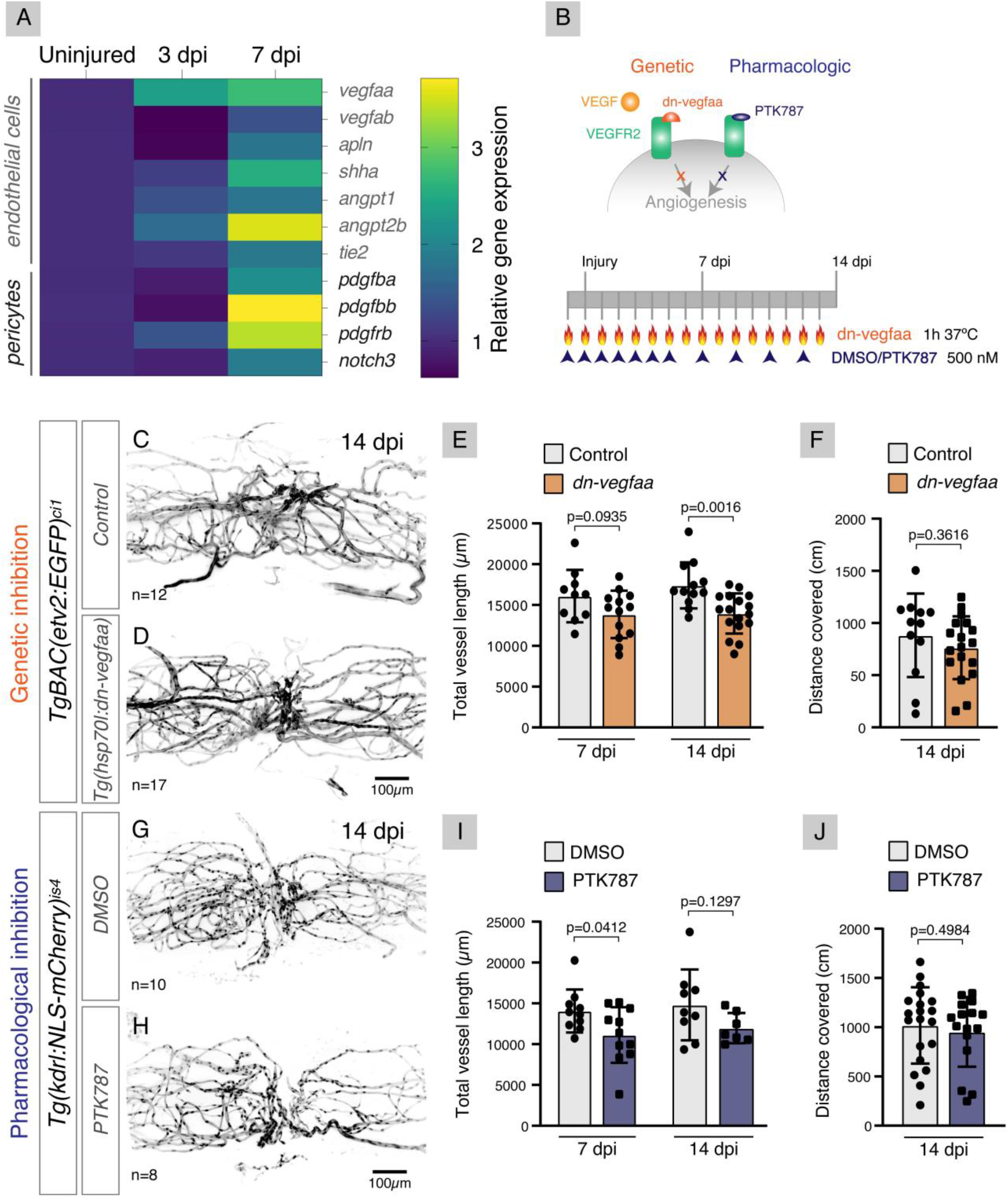
Inhibition of Vegf/Vegfr2 pathway has limited effect on vascular repair. A. Heatmap of the relative expression of genes involved in regulation of angiogenesis (grey) and pericyte recruitment (black) measured by qPCR and standardised to *gapdh*. B. Schematic of the protocols used to inhibit the Vegf/Vegfr2 pathway, either genetically (*Tg(hsp70l:dn-vegfaa)*) or pharmacologically (PTK787). C,D. Projections of light sheet images of whole spinal cords at 14 days post-injury (dpi) with contusion, with labelled endothelial cells (*TgBAC(etv2:EGFP)*^*ci1*^), in the absence (C) or presence (D) of the *dn-vegfaa* transgene after repeated heatshocks. A similar vessel pattern was observed in both conditions. E. Total vessel length at 7 and 14 dpi show a small decrease with *dn-vegfaa* when compared to Control (significantly different at 14 dpi). F. Distance covered in a open-field test at 14 dpi shows similar values between Control and *dn-vegfaa*. G,H. Projections of light sheet images of whole spinal cords at 14 dpi with labelled endothelial nuclei (*Tg(kdrl:NLS-mCherry)*^*is4*^), exposed to DMSO (G) or PTK787 (H). Both conditions show similar vascular patterns. I. Total vessel length at 7 and dpi show a non-significant decrease in the presence of PTK787 when compared to DMSO. J. Distance covered in a open-field test at 14 dpi is similar between both conditions. Data represent means ± SD. Statistical tests: Unpaired t test (E,F,I); Mann Whitney test (J).

Among the endothelial-related genes up-regulated in response to injury, *vegfaa* was of particular interest, due to its known role as a pro-angiogenic factor, including in the developing spinal cord (James et al., 2009). Moreover, *vegfaa* showed an early expression consistent with the timing of endothelial proliferation. For these reasons, we targeted Vegfaa activity to determine its role in the re-vascularisation of the injured spinal cord. We first used a genetic approach to block Vegfaa downstream signalling, based on the heat-shock-driven expression of a dominant-negative form of *vegfaa* (*Tg(hsp70l:dn-vegfaa)*) (Fig. 7B). This transgenic line was previously used to inhibit heart re-vascularisation after injury in adult zebrafish (Marín-Juez et al., 2016). To activate the *dn-vegfaa* transgene we exposed fish to daily heat-shocks from the day before SCI to the day before spinal cord collection. We confirmed by qPCR that our heat-shock protocol activated the endogenous *hsp70l* promoter and the artificial expression of *vegfaa* after repeated heat-shocks (Supp. Fig. 8A). We next analysed the vasculature (*TgBAC(etv2:EGFP)*^*ci1*^) in wholemount spinal cords at 7 dpi (not shown) and 14 dpi (Fig. 7C,D), which showed similar patterns in the absence (Control) and presence of the transgene (*dn-vegfaa*). The quantification of total vessel length in the injured region revealed only small decreases in *dn-vegfaa* fish, which were significantly different from controls at 14 dpi (Fig. 7E). Consistent with the small effect on total vessel length, endothelial proliferation at 3 dpi was also unaffected by *dn-vegfaa* overexpression (Supp. Fig. 8B,C). The reduction in vascular density observed at 14 dpi also had no effect on the motor recovery of the *dn-vegfaa* fish, as measured by the distance covered in an open-field test (Fig. 7F). These results suggest that the inhibition of Vegfaa activity is not sufficient to efficiently block spinal cord re-vascularisation.

To confirm this result we used an alternative approach to inhibit Vegf/Vegfr2 signalling, using the Vegfr inhibitor PTK787 (Fig. 7B). This compound was previously used to inhibit re-vascularisation after caudal fin amputation in adult zebrafish (Bayliss et al., 2006). The analysis of the vascular distribution at 7 dpi (not shown) and 14 dpi (Fig. 7G,H) again showed similar patterns between DMSO-treated and PTK787-treated fish. Likewise, the quantification of total vascular length showed a small decrease in PTK787-treated fish (Fig. 7I). The minimal effect of the drug on the vasculature also did not affect the swimming capacity at 14 dpi (Fig. 7J), as observed with the *dn-vegfaa* transgene.

These data support a partial role for Vegfaa signalling in vessel re-growth during spinal cord regeneration and suggest that additional pathways may be involved in promoting spinal cord re-vascularisation.

Taken together, these results reveal that the regeneration of the zebrafish spinal cord also includes the repair of the vascular system, with an early angiogenic phase (partially Vegfr2-dependent), followed by intermediate/late phases of pericyte recruitment and vessel stabilisation and remodelling (Fig. 8).

**Figure 8.**
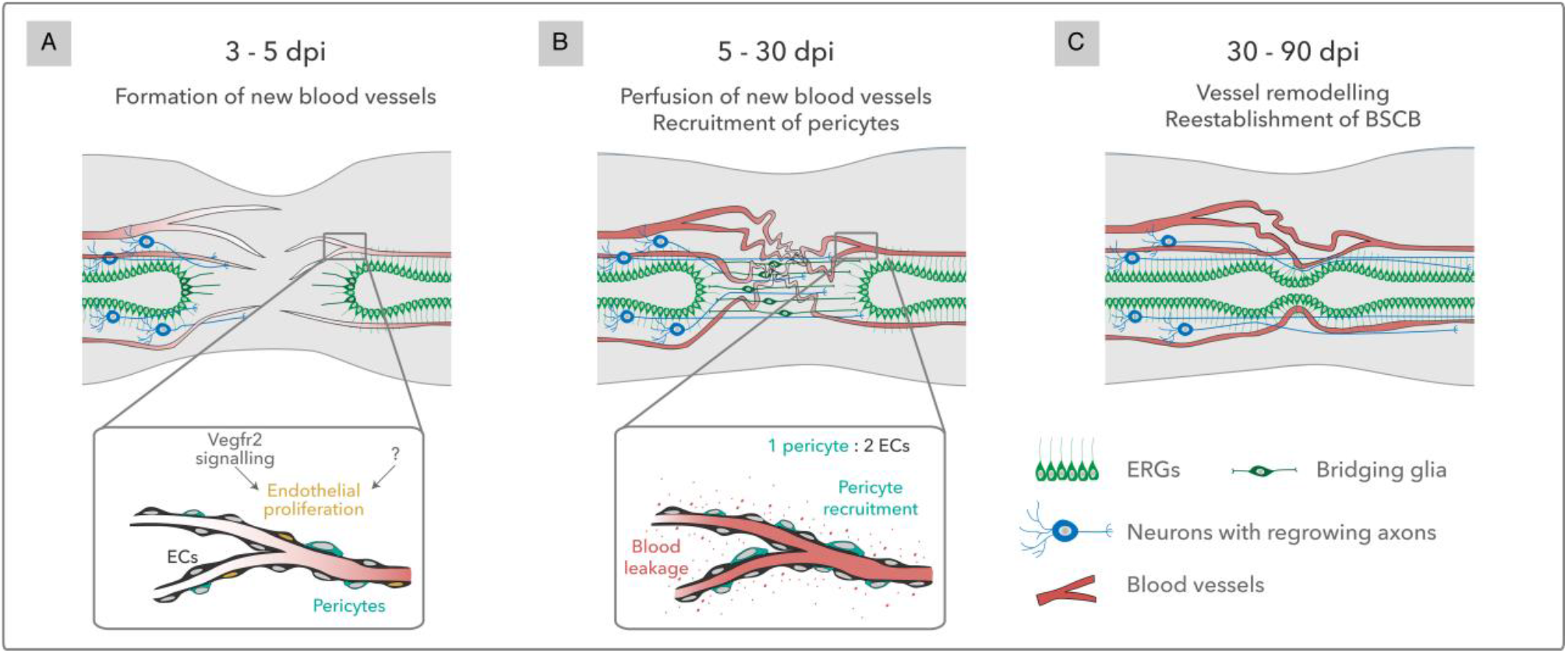
Model of vascular repair in the zebrafish spinal cord. A. SCI triggers early endothelial proliferation via Vegfr2 signalling and other unidentified pathways. New blood vessels invade the injured tissue before glial projections and regrowing axons. B. Rostral and caudal growing vessels anastomose in the injured region and become perfused around 7 dpi. The initial vascular repair creates a network of irregular vessels. Although pericytes are recruited to new blood vessels from 5 dpi, vessels remain permeable until 30 dpi. C. Between 30 and 90 dpi the vascular network is remodelled and the BSCB is reestablished, allowing for the long-term and functional vascularisation of the regenerated spinal cord.

## Discussion

The central nervous system is a metabolically demanding tissue that requires the optimised distribution of blood, while also being sensitive to several blood components, which demands a tight control of the traffic between blood and nervous cells. This balance is achieved during neurovascular development and should again be re-established after a traumatic injury if the tissue is to regain its full function. However, after SCI in mammals both vessel distribution and BSCB integrity show long-term impairments in the damaged tissue (Oudega 2012; Yao et al., 2021). Here we report the first detailed characterisation of the spinal cord vasculature in developing and adult zebrafish and show that blood vessels are successfully repaired after spinal cord injury in this regenerative organism.

The zebrafish spinal cord is vascularised in a late larval stage, with the start of endothelial migration into the spinal cord described to occur between 12 (Wild et al., 2017) and 18 dpf (Matsuoka et al., 2017). Here we propose that the size of a fish, and in particular the size of the spinal cord, determines the timing of vascular ingression, more than the fish age. Even though the ages analysed in the different studies are different, our hypothesis is supported by the fact that the average spinal cord area at the start of vessel entry is comparable between our study (8378 ± 2106 µm^2^ in 35-42 dpf fish) and the Matsuoka et al. study (8268 ± 1673 µm^2^ in 18-20 dpf fish; n=3, measured in transverse sections from figures and movies in Matsuoka et al., 2017). The age difference between studies could be explained by differences in husbandry conditions, such as type and availability of food and tank density, which can impact growth speed (Kolb et al., 2018).

The spinal cord area at which vessels start invading the spinal tissue corresponds to a 50 µm radius, which is consistent with the distance of most cells to its nearest capillaries (<100 µm) (Krogh, 1919; Place et al., 2017). Below this distance oxygen diffusion from the PNVP could be sufficient to reach the centre of the tissue and above this distance cells could start activating hypoxic signals and induce angiogenesis. Further work is needed to determine if the tissue becomes hypoxic and which cells and pathways are involved in attracting vessels into the spinal cord. Spinal neuron- and radial glia-derived signals (Sflt1 and Vegfab) were shown to affect the distribution of vessels around the spinal cord (PNVP) in earlier stages (Matsuoka et al., 2016, 2017; Wild et al., 2017) and these cell types could also be involved in promoting vessel entry in later stages.

Our findings also reveal that vessels entering and spreading into the spinal cord are accompanied by pericytes. The pericytes recruited with the invading sprouts possibly derive from the perivascular population that supports the PNVP vessels (this study and Tsata et al., 2021). Endothelial-derived Pdgfb is likely involved in attracting Pdgfrb^+^ pericytes to the nascent blood vessels. During mouse brain development PDGF-B/PDGFRß signalling is essential for pericyte migration and survival (Lindahl et al. 1997). As the brain vasculature grows, pericytes also proliferate and help regulate EC number and BSCB assembly (Lindahl et al. 1997; Hellström et al., 2001). During zebrafish spinal cord development, we also observed pericyte expansion in proportion to the increase in endothelial number, with a ratio of 1 pericyte to 3 ECs that is maintained until adulthood. Whether pericytes contribute to the establishment of the BSCB in zebrafish remains to be uncovered.

Our analysis of the spinal cord vasculature extended to the adult stage, where we described the stereotypic organization of blood vessels with larger vessels entering in dorsal and ventral positions, extending longitudinally along the central canal and spreading peripherally into the grey matter. Unlike the human spinal cord (Bosmia et al., 2015), peripheral irrigation from a vasocorona appeared to be absent in zebrafish, suggesting that a peripheral source is not necessary to irrigate the small-sized spinal cord. Ventral and dorsal ingression points were also regularly distributed along the vasculature, raising the question of how the arterial and venous systems are segregated in the zebrafish spinal cord (for example, dorsal vs ventral or in alternating segments). Further work addressing the arterial-venous identity of ingressing vessels will help answer this question and understand the circulatory route in the zebrafish spinal cord.

The adult zebrafish spinal cord has a developed BSCB, with associated pericytes and radial glial cells and tight junctions and other intercellular junctions between ECs. The presence of these components is indicative of a controlled transit of cells and molecules between blood and nervous tissue.

In this study we also provide a detailed temporal and spatial description of the vascular response to SCI. We show that the lesion severs blood vessels, but new vessels quickly reappear in the injury epicentre, from both rostral and caudal regions, and before glial projections and axonal sprouts. Endothelial proliferation contributes to the re-vascularisation of the injured tissue, but is temporally restricted to early time points, suggesting that endothelial migration may play a role afterwards. The invading vessels often travel together and create agglomerates at the injury core that appear to fuse the rostral and caudal sides of the vascular network. Short-term perfusion of a tracer dye showed that circulation is rapidly re-established in the repaired vasculature. However, the vessels initially formed have a tortuous morphology and some have a larger calibre than normal. Tortuous vessels have been described as transient in wound healing (Chong et al., 2017) and chronic in tumours (Nagy et al., 2009). In our system the tortuous vessels are transient, suggesting that the repaired vasculature is remodelled to remove the dysfunctional vessels and optimise circulation. As a result, the final vasculature has fewer and more linear vessels than pre-injury. Tortuosity can be a result of high local VEGF-A levels or the type of VEGF-A isoforms expressed (Nagy et al., 2009) and is frequently associated with hyperpermeability (Hashizume et al., 2000). Consistent with the presence of tortuous/dysfunctional vessels, vessels were permeable to a large fluorescent dextran until 30 dpi. These results revealed an unexpected delay between the formation and the maturation of new blood vessels, during which the tissue is exposed to blood components and invading immune cells. Nevertheless, by 90 dpi, when most atypical vessels had been removed, vessel permeability had also decreased to pre-injury levels, indicating that the repaired spinal tissue is supported by long-term vessels with a functional BSCB.

The mechanisms underlying the recovery of the BSCB are still unclear, but pericytes are likely involved. New blood vessels rapidly attracted pericytes, formed through proliferation and possibly migration of existing pericytes. The Pdgfrb/Pdgfb pathway is likely involved in pericyte recruitment, as we saw receptor and ligand upregulation in response to injury. Although pericytes started to be recruited early on, pericyte density only reached peak levels between 30 and 60 dpi, which corresponded to the period when vessels become less permeable. The gradual accumulation of pericytes may be important to promote the recovery of the barrier. Alternatively, it is possible that, despite being present in high density, early pericytes are unable to provide sufficient surface coverage of vessels, which can be correlated with vessel permeability (Winkler et al., 2012). Additional studies need to be performed to understand the role of pericytes in BSCB recovery, for example through regulation of endothelial transcytosis and astrocyte end feet function (Armulik et al., 2010). In addition, other BSCB modifications, such as intercellular junctions, were not investigated in this study, but it is likely that the regulation of these structures is also important for the reestablishment of the barrier.

Pericytes may also be important for the regulation of angiogenesis. Depending on the context, pericytes can have a positive effect on angiogenesis by expressing VEGF and serving as pioneer cells for growing vessels (Amselgruber et al., 1999; Reynolds et al., 2000); or a negative effect by limiting endothelial proliferation and promoting vessel maturation (Hellström et al., 2001). Pericytes were associated with angiogenic vessels at 3 dpi, consistent with a guiding role for endothelial sprouts, but were also present in high density at 5 and 7 dpi, when endothelial proliferation declines. The role of pericytes in this process will need to be addressed by inhibiting pericyte recruitment in the injured spinal cord.

The role of Vegfa signalling in spinal cord revascularisation was also examined in this study, but surprisingly revealed only a small contribution of the pathway. Although Vegfa is a prevalent pro-angiogenic signal in multiple contexts, including vessel ingression during spinal cord development (James et al., 2009), and *vegfaa* expression was detected in the injured spinal cord, inhibition of the Vegfaa pathway had only a small effect on the formation of new vessels. These results suggest that other signals are involved in the revascularisation of the spinal cord in zebrafish or can compensate for the absence of Vegfaa signalling. It is also possible that the strategies used here were inefficient in targeting the spinal cord, even if previously used to successfully inhibit vascular regrowth in zebrafish heart (*dn-vegfaa*; Marín-Juez et al., 2016) and fin (PTK787; Bayliss et al., 2006) regeneration. This explanation is not supported by the strong activation of the *hsp70l* promoter in the spinal tissue induced by our heat-shock methodology, arguing that *dn-vegfaa* is efficiently expressed but insufficient to block vascular repair. Other pro-angiogenic factors identified in this study (*shha, angpt2a*) and possible cellular sources of angiogenic signals (pericytes, ERGs, neurons, immune cells) will be targeted in the future to assess their role in this process. Vegfa inhibition also had no effect on swimming recovery, suggesting that Vegfa is not a modulator of axonal growth during spinal cord regeneration, as has been described in developmental contexts (Wälchli et al., 2015).

The inability to block vascular repair using Vegfa inhibition also prevented us from assessing the role of blood vessels during regeneration, either to provide support for the tissue or as a source of pro-regenerative signals. Blood vessels are part of the brain neural stem cell (NSC) niche and support NSC proliferation and neurogenesis (Shen et al., 2008; Tavazoie et al., 2008). We detected new vessels preferentially in the central region and, therefore, in a position to promote proliferation and differentiation of NSCs here located. Blood vessels can also support axonal regrowth during peripheral nerve regeneration, where vessels enter the connecting bridge first and serve as substrate for migrating Schwann cells and associated regrowing axons (Cattin et al., 2015). In a similar way, our study shows that blood vessels are among the first structures to enter the injury gap, before glial cells and axons, and are associated to regrowing axons, raising the possibility that vessels act as pioneers for regrowing axons. In addition, pericytes (associated or not to vessels) may also play a role in axonal guidance, for example by secreting pro-regenerative ECM components, as observed in the larval spinal cord pericytes (Tsata et al., 2021).

In conclusion, this study establishes the vascular system as a new component of the regenerative program in the zebrafish spinal cord, but further work is be needed to understand how vascular repair is regulated and how blood vessels influence other components of the injury response. This work also highlights the benefits of performing SCI studies in adult zebrafish, since larval models miss the contribution of the vascular component.

## Materials and Methods

### Fish husbandry and zebrafish lines

Zebrafish (Danio rerio) lines were raised and maintained at 28.5ºC in a 10/14h dark-light cycle, according to FELASA recommendations (Aleström et al., 2020). Fish were fed a diet of once daily dry food (Zebrafeed, Sparos) and once daily live food (Artemia). We used AB wild-type strain and transgenic lines that were previously established: *Tg(kdrl:HRAS-mCherry)*^*s896*^ (Chi et al., 2008); *TgBAC(pdgfrb:citrine)*^*s1010*^ (Vanhollebeke et al., 2015);*Tg(kdrl:NLS-mCherry)*^*is4*^ (Wang et al., 2010); *Tg(fli1:EGFP)*^*y1*^ (Lawson and Weinstein, 2002); *Tg(gfap:GFP)*^*mi2001*^ (Bernardos & Raymond, 2006); *Tg(hsp70l:vegfaa121-F17A)*^*bns100*^ (hereafter *Tg(hsp70l:dn-vegfaa*) (Marín-Juez et al., 2016); *TgBAC(etv2:EGFP)*^*ci1*^ (Proulx et al., 2010). Procedures with zebrafish were performed in compliance with animal welfare legislation and were approved by the ethical approval committee of the Portuguese veterinary department (licence reference 0421/000/000/2021, Direcção Geral de Agricultura e Veterinária).

### Spinal cord injury and motor behaviour analysis in adult zebrafish

Adult zebrafish of either sex, between 4-12 month-old, were randomly assigned to experimental groups. Fish were anaesthetised by immersion in 0.6 mM tricaine (MS222, Sigma) in system water. Fish were placed with the left side up on a sponge to make a longitudinal incision on the side of the fish, midway between the base of the skull and the dorsal fin, and the muscle fibres were separated to expose the vertebral column. For the contusion injury, the spinal cord was compressed dorsoventrally using forceps (Dumont #55, FST) (Hui et al., 2010). For the transection injury, the spinal cord was completely transected using fine scissors (#15000-03, FST) (Becker et al., 1997). The sham injury was performed by making an incision at the side of the animal but leaving the spinal cord intact. The wound was closed with forceps and the fish was placed in system tanks with 6-10 other injured fish in a ZebTEC Stand-Alone Toxicology Rack (Tecniplast).

To evaluate the motor recovery after SCI we performed an open field test at 14 dpi (Hui et al., 2010). Individual fish were placed in 9.5×16.5 cm tanks with system water with a light source underneath and allowed to acclimate for 5 minutes before recording the swimming behaviour for 5 minutes from a top view (Ikegami digital video camera). The swimming behaviour was tracked using the EthoVision XT 15.0 software (Noldus) to obtain the total distance traveled for 5 minutes. The obtained data was plotted and analysed statistically using GraphPad Prism 8.

### Heat-shock and drug treatments

For the heat-shock treatments, zebrafish were placed in pre-warmed water at 37ºC for 1 hour at a density of 6 fish per litre. Heat-shock treatments were performed once a day and fish were afterwards returned to 28.5ºC in the Toxicology Rack.

For the treatment with PTK787 (Vatalanib succinate, #5680, Tocris), up to 4 adult fish were placed in 700 ml of E3 medium (5 mM NaCl; 0.17 mM KCl; 0.33 mM CaCl2; 0.33 mM MgSO4) containing 500 nM of PTK787 (from a 10 mM stock dissolved in DMSO) or 0.005% DMSO in control fish. Treated fish were kept in the dark at 28.5ºC and the medium was changed daily and new drug/vehicle added daily or every 2 days as shown in the figure schematic.

### General image acquisition and processing details

The systems, objectives and z-stack intervals are detailed in the subsections. The pinhole aperture was adjusted to keep the same optical slice thickness between the different channels. The laser intensities were kept at the same level between samples and conditions in the same experiment. Images were processed using Fiji (Schindelin et al., 2012) and Adobe Photoshop CS4 and figures were prepared in Adobe Illustrator CS4.

### Tissue sectioning in juvenile and adult zebrafish

Fish were sacrificed in 15 mM tricaine and fixed in 4% paraformaldehyde (PFA) at 4ºC overnight. In juvenile fish the head was removed and the rest of the body was fixed; in adult fish the vertebral column was dissected and placed in fixative. After fixation the sample was washed 3 × 5 min in PBS and, for adult fish, the spinal cord was isolated from the vertebral column using forceps. The samples were equilibrated in sucrose solution (15% sucrose; 0.12M phosphate buffer (PB) pH 7.2) at 4ºC overnight, embedded in gelatin (7.5% gelatin; 15% sucrose; 0.12M PB pH 7.2) at 37ºC, frozen in isopentane cooled to -40ºC in dry ice and stored at -80ºC. In juvenile samples the trunk region was cryosectioned in 25 µm-thick transversal slices collected in 4 alternating slide sets. Adult spinal cords were cryosectioned in 14 µm-thick transversal sections mounted in 6 alternating slide sets, or in 25 µm-thick longitudinal slices collected in a single slide.

### Immunohistochemistry in sections and wholemount samples and image acquisition

To perform immunostaining in sections the gelatin was removed from the cryosections using PBS heated to 37ºC (4 × 5 min washes). After incubation with a blocking solution (1% BSA in PBS; 0.1% Triton X-100 in PBS (PBSTx)) for 1 hour at RT, the sections were incubated in primary antibody solution at 4ºC overnight. After 5 × 5 min washes in PBSTx, the sections were incubated with the secondary antibody (1:1000 in blocking solution) and 1µg/ml DAPI (#D9564, Sigma) for 2 hours at room temperature (RT). Details of the primary and secondary antibodies used are described in Supp. Tables 1 and 2. After incubation with the secondary antibodies, the sections were washed in PBS and mounted in Mowiol solution (2.4 g Mowiol; 6 g glycerol; 6 ml dH2O; 12 ml 0.2 M Tris buffer pH 8.5) with #1.5 coverslips and allowed to dry overnight. Images of transversal sections shown in Fig.1 and Fig. 2 were acquired in a Zeiss LSM 710 confocal microscope with 20X Plan-Apochromat dry objective. Images were acquired as z-stacks (4 slices 1 µm apart). Images for the quantification of number of vessels (Fig.1) were acquired in a Leica DM5000B widefield fluorescence microscope mounted with a monochrome CCD camera, using a 20x HC PLAN APO objective. Acquisition started approximately at the heart section level and for at least 600 µm in fish length (7 sections spaced 100µm) for each fish. Longitudinal images were acquired in an inverted Zeiss LSM 880 confocal microscope with 20X Plan-Apochromat dry objective. Images were acquired as z-stacks (6 slices 1 µm apart). To perform immunohistochemistry in wholemount spinal cords (Supp. Fig. 2), samples were fixed and placed in Scale A2 solution at 4ºC for at least 2 days. The spinal cords were then treated with collagenase (2 mg/ml in PBS) for 25 minutes at RT, followed by a wash in PBSTx and incubated in 50 mM glycine in PBSTx for 10 minutes at RT. Samples were then incubated in 10% blocking solution (10% goat serum; 1% DMSO; 1% Triton X-100 in PBS) at 4ºC for 2 days. The solution was then substituted with blocking solution with primary antibody and incubated at 4ºC for 5 days. The spinal cords were then washed in PBS at RT in a roller for several hours and afterwards incubated with blocking solution with secondary antibody at 4ºC for 5 days. Samples were then washed in PBS for several hours and transferred to Scale A2. Samples were stored at 4ºC until cleared and were acquired in the lighsheet microscope as described in the clearing subsection.

### Tissue clearing in juvenile and adult zebrafish and image acquisition

Samples for tissue clearing were fixed as for tissue sectioning. After washing 3×5 minutes with PBS, the samples were incubated with Scale A2 solution (4 M urea; 10% glycerol; 0.1% Triton X-100; Hama et al., 2011) at 4ºC, protected from light, until cleared (minimum of 2 weeks). For confocal imaging, samples were kept in Scale A2 and placed in a glass-bottom Petri dish (FluoroDish, #FD35COL, WPI) with a minimum of liquid. Samples were imaged in an inverted Zeiss LSM 880 confocal microscope with 25X LCI Plan-Neofluar Corr DIC water immersion objective. Images were acquired as z-stacks with slices 2.751 µm apart with the maximum depth possible (<100 µm). For light sheet imaging, samples were transferred to Scale S4 (40% D-(-)-sorbitol; 4 M urea; 10% glycerol; 0.2% Triton; 15% Dimethylsulfoxide; Hama et al., 2015) the day before imaging and kept at 4ºC protected from light. Samples were imaged in a Zeiss Lightsheet Z1 microscope with a 20X Clr Plan-Neofluar Corr nd=1.45 85mm clearing immersion objective. During imaging the spinal cord was suspended in clearing solution from a glass capillary (size 1 / ∼0.68 mm). Dual-side images of injured or control regions were acquired as z-stacks with slices 1.14 µm apart along the whole depth of the spinal cord. Dual-side stacks were fused after acquisition using ZEN 2014 SP1 software.

### Dextran Alexa Fluor 647 heart injection and detection

To evaluate vessel perfusion and permeability properties we injected the tracer dye Dextran, Alexa F uor™ 647; 10,000 MW, Anionic, Fixable (#D22914, Invitrogen). Either *Tg(kdrl:HRAS-mCherry)*^*s896*^ or *Tg(kdrl:NLS-mCherry)*^*is4*^ lines were used for the injection. The dextran-A647 solution (50 mg/ml in water) was uploaded to a glass microinjection needle using a suction tube with a mouthpiece. Fish were anaesthetised by immersion in 0.6 mM tricaine and placed with the ventral side up on a sponge soaked with tricaine solution. An incision was performed at the level of the heart. The heart was injected with 0.3-0.5 µl with the glass needle, using a stereoscope to visually control the entry of the dye in the heart. For the short-term and long-term perfusions, fish were either kept in the sponge for 1 minute or returned to system water for 30 minutes respectively, before sacrificing the fish and collecting the vertebral column. Spinal cords fixed and sectioned longitudinally (25 µm-thick) as described above. To mount the slides, gelatin was removed from cryosections using PBS heated to 37ºC (4 × 5 minutes washes). The slides were then incubated with DAPI for 20 minutes at RT and mounted in Mowiol. Imaging was performed in an inverted Zeiss LSM 880 confocal microscope with 20X Plan-Apochromat dry objective. Images were acquired as z-stacks (6 slices 1 µm apart) taken at mid-section depth. The rostral and caudal sides of the injury and a region 3 mm caudal to the injury were imaged in 3 adjacent sections that included or were near the central canal.

### EdU incorporation and detection

To label proliferating cells we used the thymidine analog EdU (5-ethynyl-2’-deoxyuridine) that is incorporated into DNA during active DNA synthesis. The EdU solution (2.5 mg/ml dissolved in PBS) was injected intraperitoneally (volume 20-40 µl adjusted to the size of the fish) using a 0.5 ml U-100 insulin syringe with 30G needle (#324825, BD Micro-Fine). EdU injections were performed in the time points described in the figure schematics. Spinal cord samples were collected, fixed and sectioned longitudinally (25 µm-thick) as described above. For EdU detection, gelatin was removed from cryosections using PBS heated to 37ºC (4 × 5 minutes washes), followed by incubation with 3% blocking solution (3% Bovine Serum Albumin (BSA); 0.5% Triton X-100 in PBS) for 30 minutes at RT. The Click-it reaction cocktail was prepared as described in the kit instructions (Click-iT™ EdU Cell Proliferation Kit for Imaging, Alexa F uor™ 647 dye, #C10340, Invitrogen), but with half the concentration of Alexa-647 dye. The slides were covered with 150-200 µl of reaction solution and incubated for 30 minutes in the dark. After 3×5 minute washes in 3% blocking solution, the slides were stained with DAPI, mounted and imaged as described for dextran-A647 detection.

### Transmission Electron Microscopy (TEM)

The vertebral column of adult zebrafish was dissected and the spinal cord isolated from the vertebrae before fixation overnight at 4ºC in 0.1 M sodium cacodylate buffer, pH 7.4, containing 2,5% (v/v) glutaraldehyde. Following 1 hour post-fixation in 1% (aq.) osmium tetroxide and 30 minutes contrast in block in 1% (aq.) uranyl acetate. Dehydration was made using ethanol gradient (50-70-95-100%) and infiltration was aided using propylene oxide and a mixture (1:1) of propylene oxide and EPON 812 resin (EMS). Samples were embedded in EPON resin and hardened at 60ºC for 36h. Sections were obtained using an ultramicrotome Reichert Supernova (Leica microsystems), semi-thin sections (500 nm) were stained with toluidine blue for light microscope evaluation. Ultra-thin sections (70 nm) were collected in Formvar (AGAR scientific) coated copper slot grids, and counter-stained with uranyl acetate and lead citrate (Reynold recipe) and screened in a Hitachi H-7650 transmission electron microscope at 100kV acceleration.

### Quantitative PCR

For injury-induced qPCR analysis, total RNA was extracted from twenty pooled 4 mm spinal cord fragments including the injury site, while for heatshock-induced expression analysis, three whole spinal cords were pooled 2 hours after the last heatshock. RNA extraction was performed using TRIzol and RNeasy Mini Kit (Qiagen #74104) according to the manufacturer’s instructions. The purified RNAs were reverse transcribed with the iScriptTM cDNA Synthesis Kit (Bio-Rad) and qPCR was performed with Power SYBR Green PCR Master mix (Applied Biosystems) and a 7500 Fast Real-time PCR system (Applied Biosystems). All experiments were completed using biological triplicates, expression levels were normalised to *gapdh* or *eef1a1* as control and the relative expression values were obtained using the CT method. Primer sequences are noted in Supp. Table 3. Normalised values were plotted using GraphPad Prism 8.

### Quantification and statistical analysis

Quantified data were organised and processed using Numbers spreadsheet (Apple) and RStudio (Version 1.4.1717). Data plotting was performed in RStudio (ggplot2 package v3.3.5) or GraphPad Prism 8 (GraphPad Software, Inc.). Plotted data were presented as mean ± standard deviation (SD). Statistical analysis was performed using GraphPad Prism 8, unless stated otherwise. Data were tested for normal distribution using Shapiro-Wilk normality test and parametric (One-way ANOVA followed by Dunnett’s multiple comparisons test; Unpaired t-test) and non-parametric tests (Kruskal-Wallis test followed by Dunn’s multiple comparisons post-hoc test; Wilcoxon matched-pairs signed rank test; Mann Whitney test) were used accordingly. All P-values were included in the graphs, with the exception of graphs with multiple comparison tests between all conditions, in which only P-values <0.05 were shown.

#### Quantification of number of vessels in juvenile fish

Images of transversal sections acquired with the DM5000b microscope were scaled and the fluorescent channels merged using Fiji. In each image the spinal cord was measured for area, dorsal-ventral and left-right lengths using Fiji. For each image, the spinal cord limits were defined and blood vessels inside were counted manually. The obtained spinal cord measurements, blood vessels or fish lengths were plotted and analysed using ggplot2.

#### Kugler analysis of vessels in wholemount spinal cords

Data analysis of light sheet images was performed in Fiji, using an adapted 3D image analysis workflow developed by Kugler et al., 2022. Quantification of vessel branch length and position, thickness and tortuosity was done as described by the authors, with the following changes to the macro: manual regions of interest (ROIs) with 500×500x[z size] µm were defined for each spinal cord image using Fiji’s polygon tool and the threshold range for image segmentation was tested and adapted to our data set. To have additional information about the maximum blood vessel thickness, a Fiji plug-in BoneJ – Thickness was integrated in the workflow, using segmented images from the workflow as input. Quantification of vessel length in *Tg(hsp70l:dn-vegfaa)* and PTK787 and control samples was performed blinded. Total vessel length, branch tortuosity index (tortuosity = branch_length / euclidean_distance) and maximum thickness were plotted and analysed statistically using GraphPad Prism 8.

The heatmaps of vessel length per bin were created using ggplot2 (geom_tile function). The middle xy position of each vessel branch was calculated and binned in 100×100 µm bins in a 500×500 µm region at the site of the injury. The sum vessel length in each bin was then calculated for each sample and plotted as a heatmap. A similar approach was used to bin vessels according to rostral-injury-caudal position (200-100-200 µm bins along the RC axis) and dorsal-middle-ventral position (200-100-200 µm bins along the DV axis). To quantify the central-lateral positions the middle z positions of branches were normalised to the total z size of each spinal cord and binned according to lateral-central-lateral position (25-50-25% bins). The total vessel length in each bin for each sample and timepoint was then normalised to the average vessel length in each bin in the uninjured condition to obtain the fold change and the average normalised values per region and timepoint were plotted as a heatmap using GraphPad Prism 8.

#### Quantification of number of endothelial nuclei in light sheet images of wholemount spinal cords

The number and position of *kdrl:NLS-mCherry*-positive nuclei per spinal cord was quantified manually using the Cell Counter plugin available in Fiji, from maximum projections of light sheet images of whole spinal cords (500×500x[z size] µm). To obtain the fold change from 1 dpi, the number of endothelial nuclei for each sample was normalised to the average nuclei number in the 1 dpi condition. The obtained data was plotted using ggplot2 and analysed statistically using GraphPad Prism 8. Heatmaps of nuclei number and position were created using ggplot2 as described for total vessel length.

#### Quantification of extravasated dextran-A647 in spinal cord sections

The intensity levels of dextran-A647 were quantified in six 50×50×5 µm regions of interest (ROIs) in maximum projection images, positioned in the extravascular tissue and excluding the central canal (Supp. Fig. 5A-B). The average intensity between ROIs in each image was then averaged between the 3 histological sections for each sample. The obtained data was plotted and analysed statistically using GraphPad Prism 8.

#### Quantification of proliferating ECs in spinal cord sections

The number of *kdrl:NLS-mCherry* positive and *kdrl:NLS-mCherry/*EdU double positive nuclei per ROI were quantified manually using the Cell Counter plugin available in Fiji, from maximum projection images of longitudinal sections (850.2×425.1×5 µm region centred in the injury). The proportion of proliferating ECs was calculated as the fraction of *kdrl:NLS-mCherry*/EdU double positive nuclei over the total number of *kdrl:NLS-mCherry* nuclei. The absolute or relative values for each ROI were then averaged between the 3 histological sections for each sample. The obtained data was plotted and analysed statistically using GraphPad Prism 8.

#### Quantification of number of ECs and pericytes in confocal images of wholemount spinal cords

The number of *kdrl:NLS-mCherry* nuclei and *pdgfrb:citrine* cells was quantified manually in maximum projections of confocal images of the left region of cleared spinal cords (340.08×340.08×90 µm depth) using the Cell Counter plugin available in Fiji. The number of ECs was normalised to the number of pericytes to obtain the EC/pericyte ratio. The obtained data was plotted and analysed statistically using GraphPad Prism 8.

## Acknowledgements

We thank the support given by Lara Carvalho and Aida Barros from the Fish Facility, the Histology and Comparative Pathology Laboratory and the Bioimaging Unit of the Instituto de Medicina Molecular - João Lobo Antunes (iMM-JLA). We also thank the Electron Microscopy Facility at the Instituto Gulbenkian de Ciência for the use of the Transmission Electron Microscope and Andreia Pinto for TEM data. We further thank Didier Stainier for the following transgenic lines: *TgBAC(pdgfrb:citrine)*^*s1010*^*;Tg(kdrl:NLS-mCherry)*^*is4*^; *Tg(hsp70l:dn-vegfaa); TgBAC(etv2:EGFP)*^*ci1*^.

## Author contributions

A.R. and L.S. conceptualised the study; A.R. performed animal and image acquisition experiments and analysed the data; M.R.C. performed the 3D vessel analysis; M.R.C. and C.d.S.-T. performed qPCR experiments; E.C.R. performed vessel quantification in juvenile fish; R.Q. performed cryosectioning; T.M. and S.C.R.S. participated in optimisation of protocols; A.R. wrote initial draft. A.R., L.S., M.R.C., C.d.S.-T. reviewed and edited the manuscript.

