## Supplemental material for "New functional vessels form after spinal cord injury in zebrafish"

#### Supplementary Figure 1

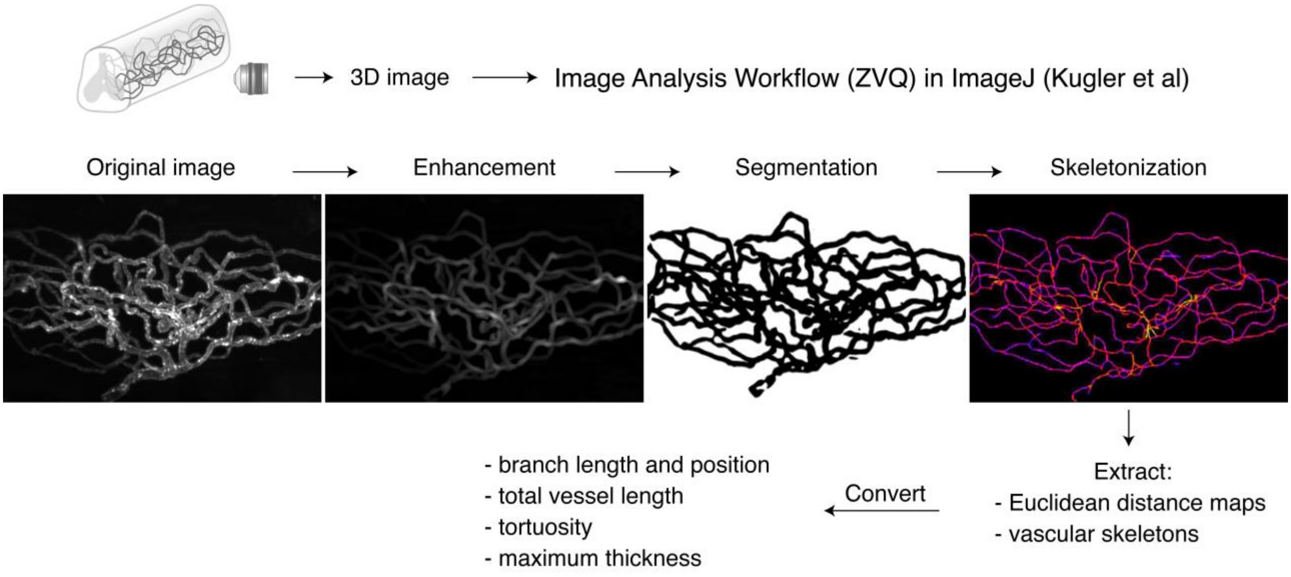


Supplementary Figure 1 - Image analysis of vessel distribution in wholemount spinal cords. Schematic of the image analysis workflow using the ImageJ-based approach developed by Kugler et al (REF) - Zebrafish Vascular Quantification (ZVQ) workflow. Cleared spinal cords with fluorescently labelled vessels were imaged using light sheet microscopy to obtain 3D images of the whole spinal cord. The 3D images were processed and quantified in ImageJ using a simplified version of the ZVQ workflow adapted for the adult zebrafish spinal cord. Images were enhanced, segmented and skeletonized to obtain Euclidean distance maps and vascular skeletons, which were then converted to obtain the length and position of individual vascular branches, total length of the vascular network, the average tortuosity and the maximum thickness.

#### Supplementary Figure 2

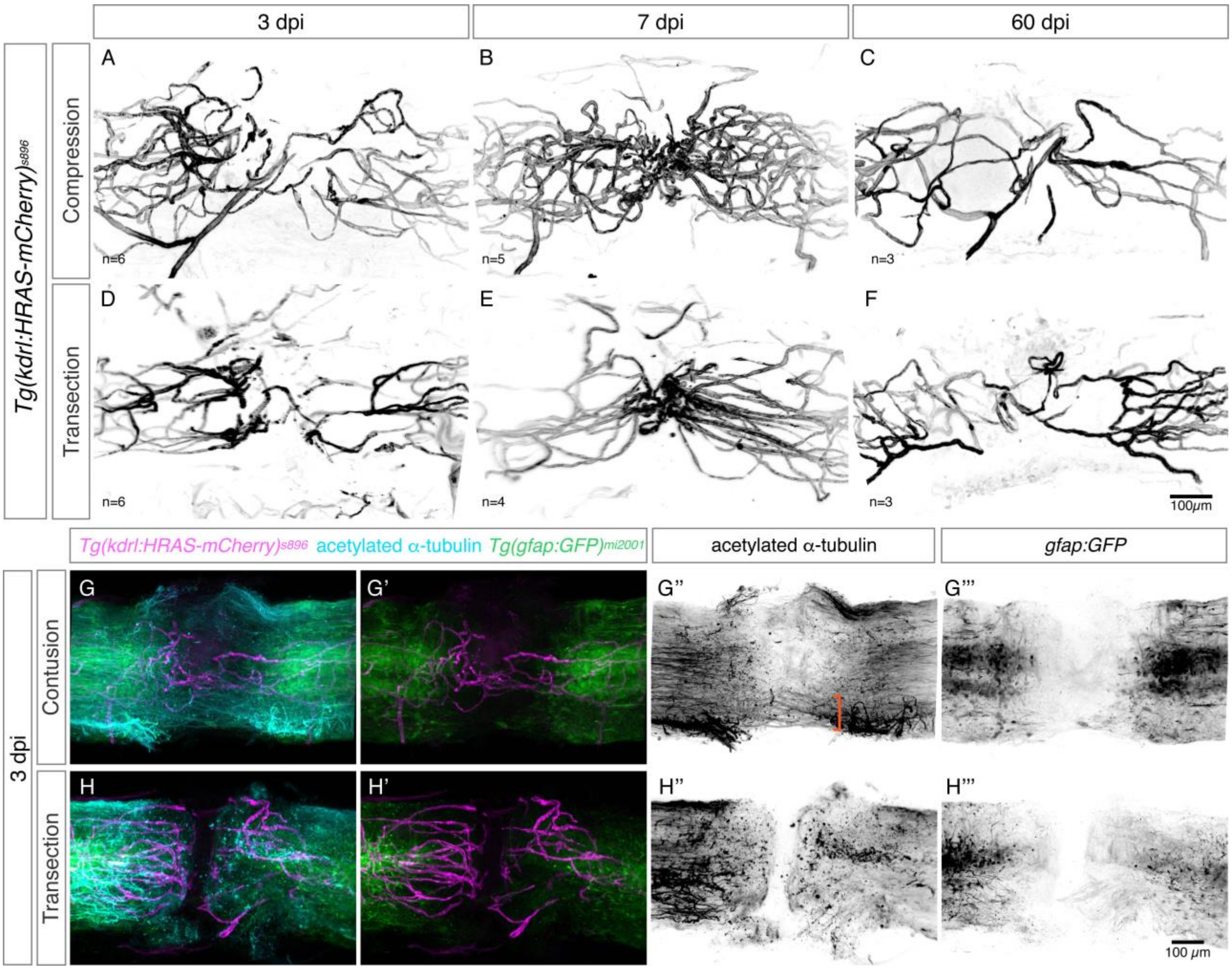


**Supplementary Figure 2 - Characterisation of the axonal and vascular response in two spinal cord injury models - contusion and transection.**

**A-F.** Projections of light sheet microscopy images of whole spinal cords with labelled endothelial cells (*Tg(kdrl:ras-mCherry)*$^{s896}$) in two models of spinal cord injury - contusion (**A-C**) and transection (**D-F**) - at 3, 7 and 60 days post-injury (dpi). A similar pattern of vascular repair is observed in both models. **G-H'''**. Projections of light sheet microscopy images of whole spinal cords with labelled endothelial cells (*Tg(kdrl:ras-mCherry)*$^{s896}$), glial cells (*Tg(gfap:GFP)*$^{mi2001}$) and axons (antibody against acetylated alpha-tubulin) in two models of spinal cord injury - contusion (**G-G'''**) and transection (**H-H'''**) - at 3 days post-injury (dpi). In a contusion model some axonal tracts remain at the injury site (orange bracket in **G''**).

### Supplementary Figure 3

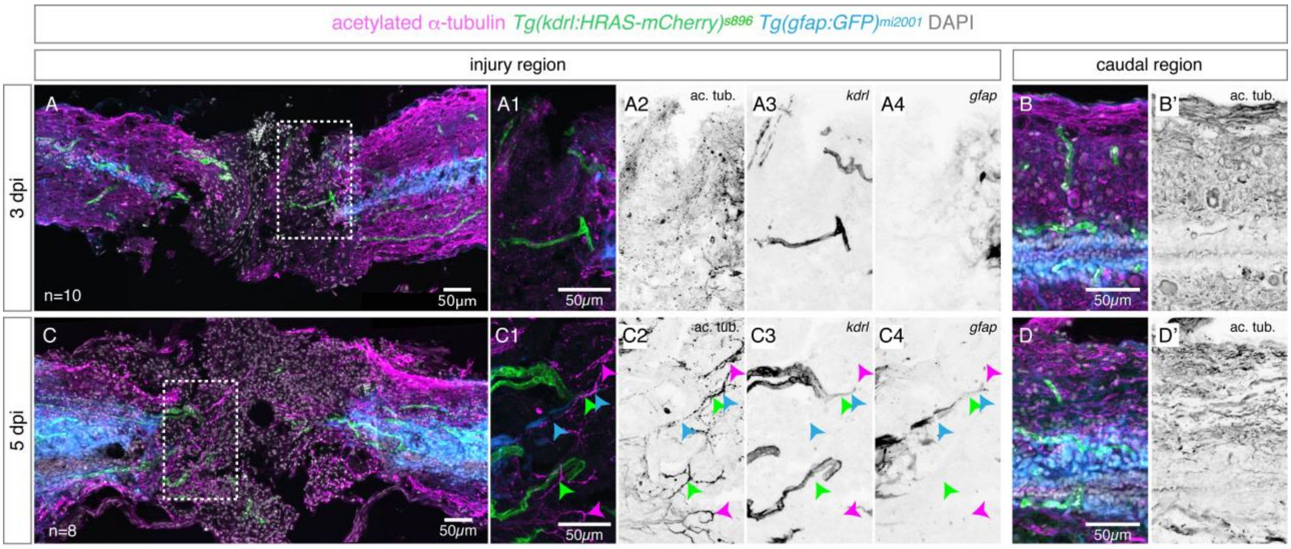


Supplementary Figure 3 - Dynamics of axonal, vascular and glial repair in the injured tissue. **A,C.** Confocal projections of longitudinal sections of transected spinal cords with labelled axons (immunostained against acetylated alpha-tubulin), endothelial cells (*Tg(kdrl:ras-mCherry)*$^{s896}$), glial cells (*Tg(gfap:GFP)*$^{mi2001}$) and nuclear DAPI staining, at 3 (**A**) and 5 (**C**) days post-injury (dpi). **A1-A4.** Magnifications of box in **A** showing blood vessels, but not axons or glial cells, present in the injured region at 3 dpi. **C1-C4.** Magnifications of box in **C** showing entry of axons, blood vessels and glial projections into the injury core at 5 dpi. Growing axons are observed adjacent to blood vessels (green arrowheads), glial projections (blue arrowheads) or alone (magenta arrowheads). **B,B',D,D'.** Region of the spinal cord 2 mm caudally to the injury.

#### Supplementary Figure 4

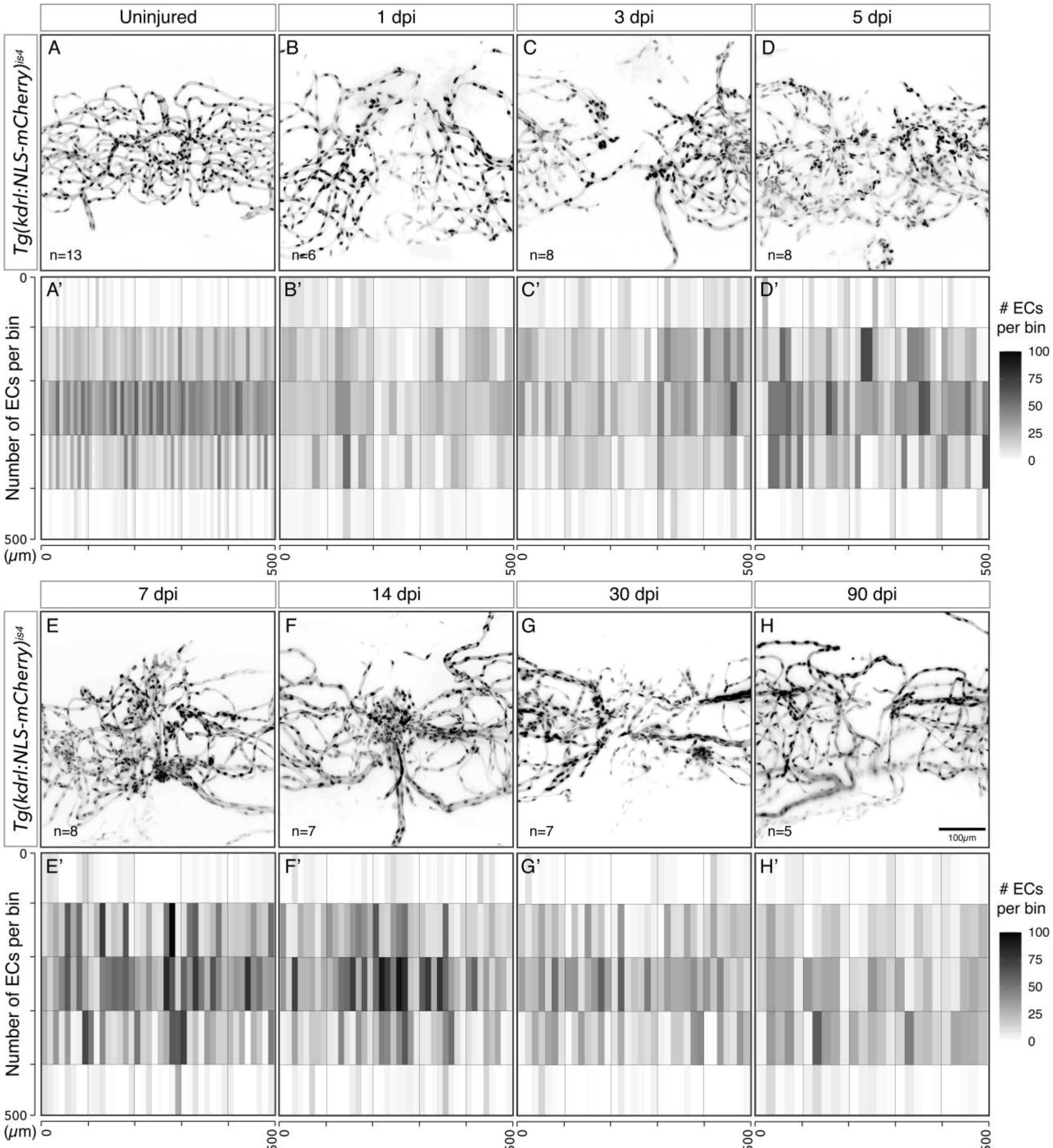


Supplementary Figure 4 - Dynamics of endothelial cell number after spinal cord contusion injury. **A-H.** Projections of light sheet microscopy images of whole spinal cords with labelled endothelial nuclei (*Tg(kdrl:NLS-mCherry)*$^{is4}$), without injury and 1, 3, 5, 7, 14 30 and 90 days post-injury (dpi) with contusion. **A'-H'**. Heatmap of the total number of endothelial cells (ECs) per 100 μm bins along the rostro-caudal and dorso-ventral axis in a 500x500 μm region. Each rectangle inside a bin corresponds to an individual sample (n=5-13).

#### Supplementary Figure 5

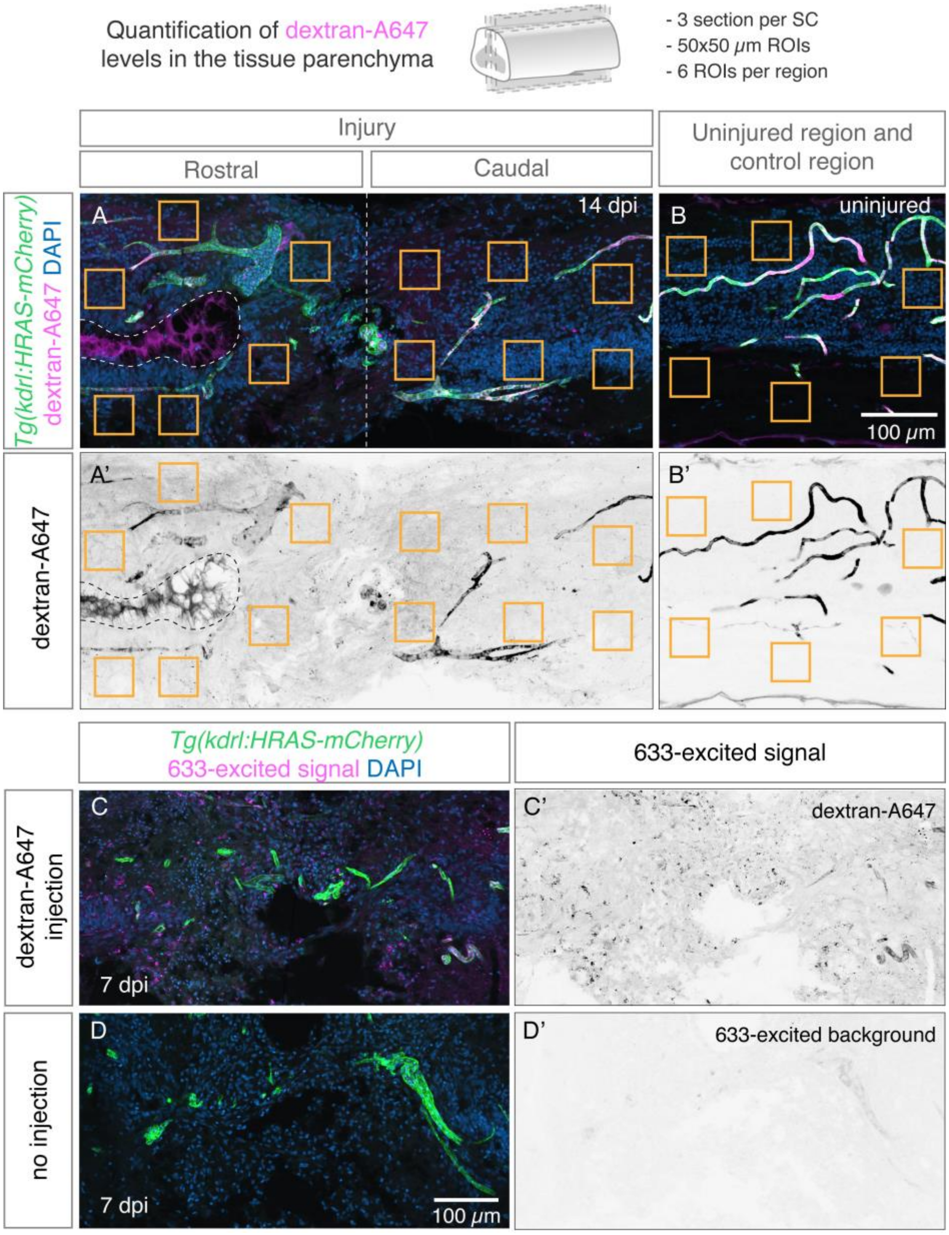


**Supplementary Figure 5 - Image analysis of vessel permeability.**

**A-B'**. Strategy to quantify dextran-A647 intensity levels in the injured spinal cords (rostral and caudal sides) (**A**,**A'**) or in control regions (**B**,**B'**), either in uninjured spinal cords or in a region 3 mm caudal to the injury. Six 50x50 µm regions of interest (ROIs) were defined in each area in the tissue outside vessels, identified by the mCherry signal. The central canal lumen (delimitated by the dashed lines in **A**,**A'**) was avoided, as the dye accumulated in this structure. **C-D'**. Confocal projections of spinal cord longitudinal sections with labelled endothelial cells (*Tg(kdrl:ras-mCherry)*$^{s896}$) and DAPI-stained nuclei at 7 days post-injury (dpi), after a 30 minute perfusion with dextran-A647 (**C**,**C'**) or without any dye injection (**D**,**D'**). After dextran-A647 injection the dye is excited by the laser 633 and the signal is detected in much higher levels in the tissue parenchyma (**C'**) when compared to 633-excited background levels in the uninjected spinal cord (**D'**).

### Supplementary Figure 6

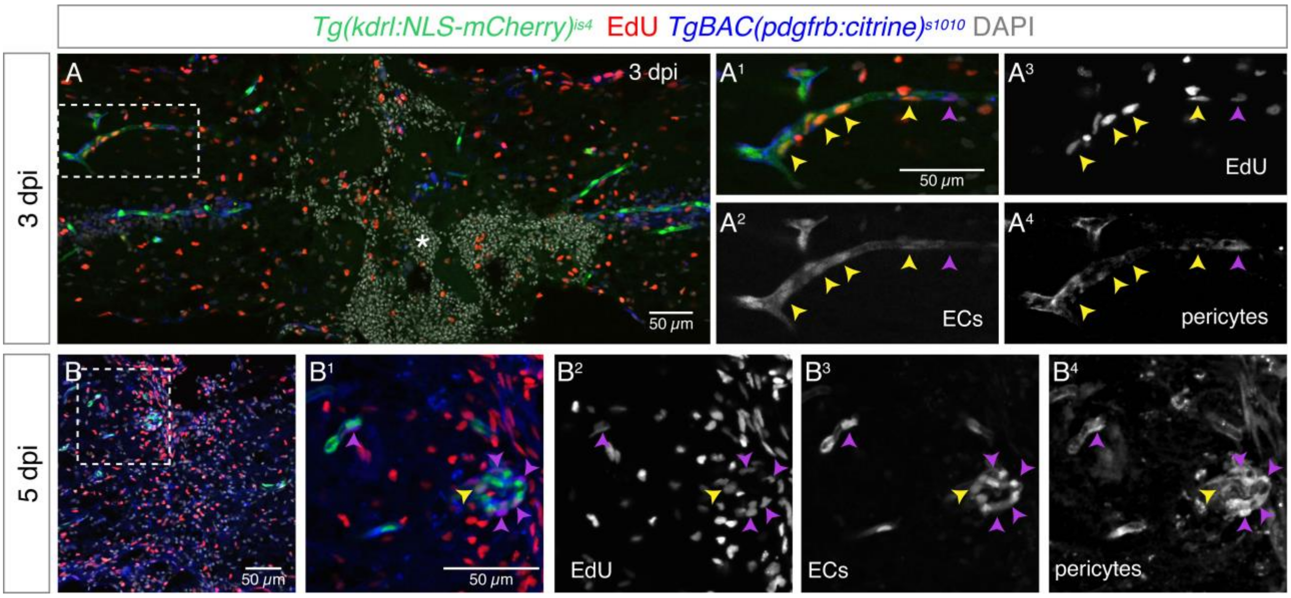


**Supplementary Figure 6 - Pericytes proliferate in response to injury.**

**A,B.** Confocal projection of spinal cord longitudinal sections 3 (A) and 5 (B) days after spinal cord injury, with EdU staining in recently divided cells, labelled endothelial cells (*Tg(kdrl:NLS-mCherry)*$^{is4}$), pericytes (*TgBAC(pdgfrb:citrine)*$^{s1010}$) and nuclear DAPI staining. EdU injection was performed 1 day before spinal cord collection. In (A) the injury epicentre is labelled by an asterisk, where a hemorrhage is still present. Magnified views ($A^1$-$A^4$,$B^1$-$B^4$) show proliferating ECs (yellow arrowheads) and pericytes (purple arrowheads).

#### Supplementary Figure 7

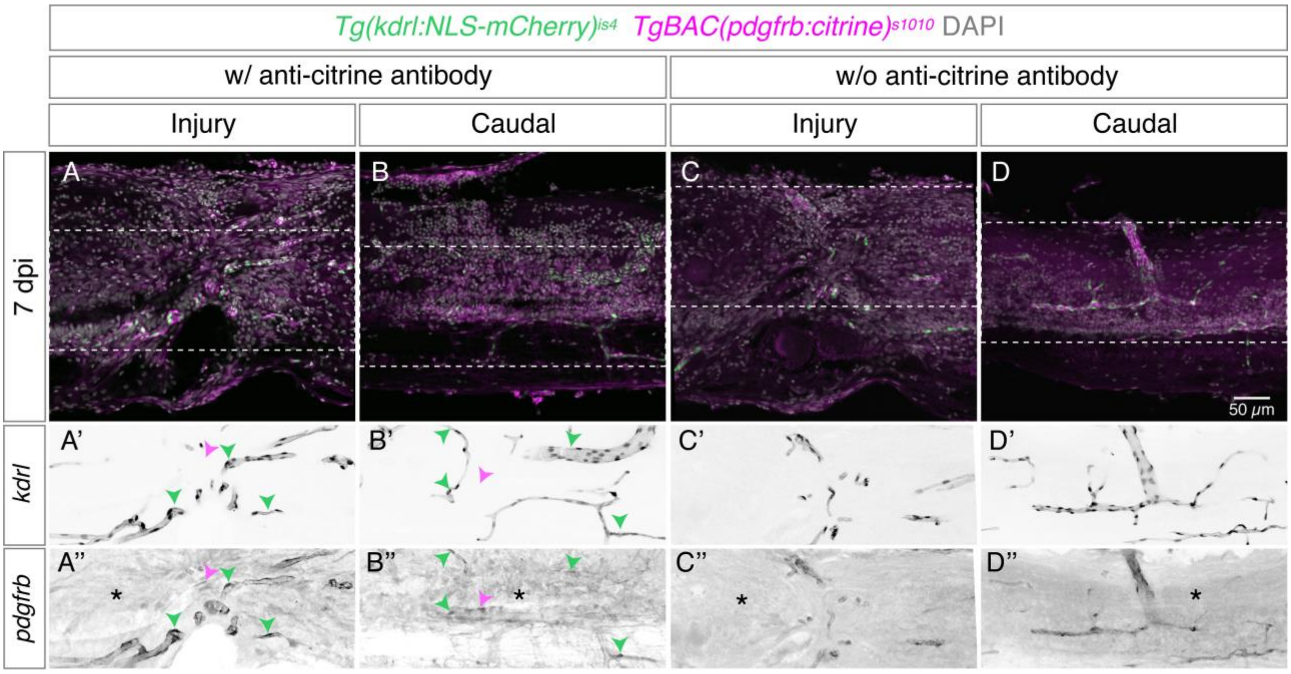


**Supplementary Figure 7 - Pericytes remain associated to blood vessels post-injury.**
**A-D.** Confocal projection of longitudinal sections of spinal cords with labelled endothelial cells (*Tg(kdrl:ras-mCherry)*$^{s896}$), pericytes (*TgBAC(pdgfrb:citrine)*$^{s1010}$) and with DAPI-stained nuclei after 7 days post-injury (dpi) with contusion. Adjacent sections labelled with (**A**,**B**) or without (**C**,**D**) an antibody against citrine were imaged in the injury region (**A**,**C**) and in a region 2 mm caudally to the injury (**B**,**D**). The pdgfrb:citrine signal was stronger with the antibody, but a strong background (asterisk) was detected in the tissue with or without the antibody staining. **A'-D'**. Inverted image of the kdrl:mCherry channel showing blood vessels. **A''-D''**. Inverted image of the pdgfrb:citrine channel showing pericytes coating blood vessels. The strongest signal is detected in cells associated with vessels (green arrowheads), but other cells in the tissue could also be positive for the transgene (magenta arrowheads), both in the injury and in a control region (**A''**,**B''**). However, these cells are masked by the high background.

#### Supplementary Figure 8

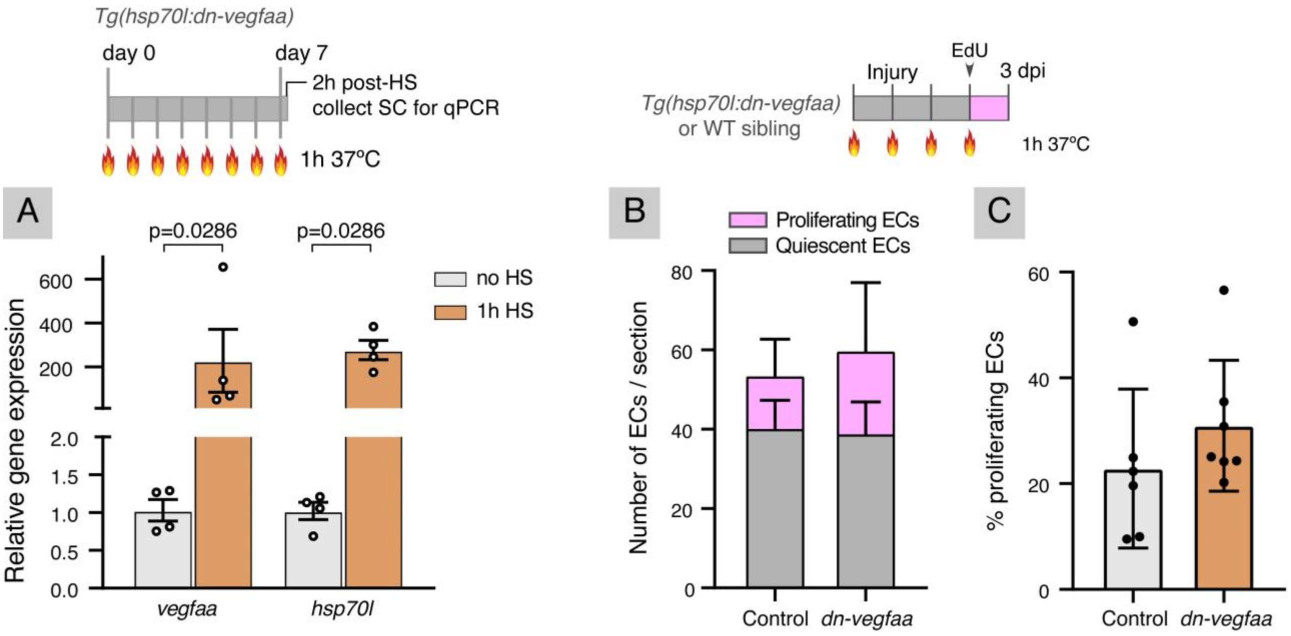


**Supplementary Figure 8 - Activation of heat-shock promoter and effect of dn-vegfaa expression on endothelial proliferation.**

**A.** Relative expression of *vegfaa* and *hsp70l* in response to heat-shock (HS), measured by qPCR and standardised to *eef1a1*. Transgenic fish *Tg(hsp70l:dn-vegfaa)* were exposed to 1h HS at 37ºC or no HS. Both *vegfaa* and *hsp70l* were significantly upregulated in response to HS. Measured *vegfaa* levels include both endogenous and transgene expression. **B.** Total number of endothelial cells (ECs) per section, grouped as proliferating ($EdU^+$) or quiescent ($EdU^-$) cells in 3 dpi wild-type controls (n=6) or *Tg(hsp70l:dn-vegfaa)* (n=7) fish exposed to daily heat-shocks from 1 day before injury to 2 dpi and injected with EdU the day before spinal cord collection. **C.** Fraction of proliferating ECs ($EdU^+$ ECs / total ECs) per section. Data represent means ± SD. Statistical test: Mann Whitney test (A).

Supplementary Table 1 - Primary antibodies

| Antigen | Host | Dilution | Source/Reference |
|---|---|---|---|
| GFP | Rabbit | 1:2000 | Abcam/ab290 |
| GFP and Citrine | Rabbit | 1:500 | Torres Pines/TP401 |
| Acetylated α-tubulin | Mouse | 1:500 | Sigma/T7451 |
| GFAP | Mouse | 1:100 | ZIRC/zrf-1 |
| HuC/D | Mouse | 1:500 | Life Technologies/A21271 |
| ZO-1 | Mouse | 1:500 | ThermoFisher Scientific/339194 |

Supplementary Table 2 - Secondary antibodies

| Against | Host | Fluorophore | Source/Reference |
|---|---|---|---|
| Rabbit IgG | Goat | Alexa Fluor 488 | ThermoFisher Scientific/A11008 |
| Mouse IgG | Goat | Alexa Fluor 633 | ThermoFisher Scientific/A21050 |
| Mouse IgG | Goat | Alexa Fluor 594 | ThermoFisher Scientific/A11005 |

Supplementary Table 3 - qPCR primer sequences

| **Gene** | **Primer Forward (5'-3')** | **Primer Reverse (5'-3')** |
|---|---|---|
| *vegfaa* | GTG CAG GAT GCT GTA ATG ATG AGG | AAT TAT GCT GCG ATA CGC GTT G |
| *vegfab* | GGT GCT GCA ATG ATG AAA TG | AAT GTC ACC CTG ATG ACG AAG |
| *hsp70l* | TGT CCA TCC TGA CCA TTG AA | CAC AAA GTG GTT CAC CAT GC |
| *angpt1* | GGC TTC CAG AAA AAC GGA GT | AAA TCC ACA GTG CCG TCT TC |
| *angpt2b* | CTG CTG GCT ATG GTC ACT CA | TCC CAT TTT CTG TCA CTC CA |
| *apln* | TGA AGA TCT TGA CGC TGG TG | GCA AAG GAG TCC TCA TGC TT |
| *notch3* | TGC CTT GTT GAA GAA TGG TG | TCC CTA TTG GCA AAG TGA GC |
| *pdgfba* | TCA CCA ACA CCA CCA ACA TC | GGG AGG AGC TTT CAT CAC TG |
| *pdgfbb* | CGC AGT CTT CTA GAC GCT CA | AGG CCA CAA CAG GAA ATC TG |
| *pdgfrb* | GAA AAG AGG ATA CCG CAT GG | GCA CCA GAA AGG AGA ACT CG |
| *shha* | AAG CCC ACA TTC ATT GCT CT | GTC CTT CAC GGC CTT CTG T |
| *tie2* | ATT ACG CCT CTC AGG ACG AC | ACT CGA TGG CCA GAT ACA GG |
| *gapdh* | GAT ACA CGG AGC ACC AGG TT | CCA GCT TGA CAA AGT GAT CG |
| *eef1a1* | TGA TCT ACA AAT GCG GTG GA | TTT GCT GGT CTC GAA TTT CC |
